# Cannabis THC:CBD Composition Affects Oligodendrocyte Progenitor Cell Characteristics Following Acute Cannabis Vapor Inhalation in Adult Male and Female Mice

**DOI:** 10.64898/2026.09.18.752658

**Authors:** Colin J. Murray, Hayley H. A. Thorpe, Hakan Kayir, Emiko Osborne, Sean W. Foster, Sophia Loewen, Mika Rogers, Haley A. Vecchiarelli, Jibran Y. Khokhar, Marie-Ève Tremblay

## Abstract

Cannabis is one of the most widely consumed substances in the world. Consumers seek out cannabis cultivars with varying levels of phytocannabinoids, primarily delta-9-tetrahydrocannabinol (THC) and cannabidiol (CBD). The effects of THC, CBD or the combination of THC:CBD have distinct outcomes on cognitive processes, cellular functions, and phytocannabinoid pharmacokinetics. The majority of research on cannabis effects on the brain has focussed on neurons, and few studies have investigated the impact of different cannabis cultivars on glia. In particular, the impact of varying levels of THC:CBD on oligodendrocyte lineage cells, which play numerous support roles in the brain essential to proper neuronal communication, is relatively unknown. This study set out to examine the acute impact of different cultivars of vaporized cannabis on oligodendrocyte lineage cells in the forceps minor of adult male and female mice. Mice were exposed to vapor from cannabis flower high in THC, high in CBD or balanced in THC:CBD over 15 minutes, and brains were fixed 30 minutes post-cannabis onset. Using immunofluorescence microscopy, we observed significant changes to oligodendrocyte progenitor cell (OPC) morphology in mice exposed to balanced cannabis, and, using correlative light and electron microscopy, we observed alterations to OPC mitochondria. The alterations observed (i.e., enlarged soma and nucleus volume, reduced density and increased area of mitochondria in the soma) in OPCs due to balanced cannabis are reminiscent of the very early changes seen during OPC differentiation. This study highlights the differing effects of cannabis cultivars on OPCs and the rapidity of the OPC response to inhaled phytocannabinoids.

**Main Points:**

- Cannabis cultivars have sex-dependent effects
- OPCs respond to vaporized cannabis within 30 minutes
- Balanced cannabis effects OPC soma size and mitochondria in male mice, reminiscent of the early changes during OPC differentiation

## Introduction

Phytocannabinoids are terpenophenolic compounds abundant in the plants belonging to the genus *Cannabis*. The two most typically abundant phytocannabinoids found in cannabis are delta-9-tetrahydrocannabinol (THC), the main intoxicating phytocannabinoid, and cannabidiol (CBD), which is generally considered not intoxicating (Pertwee, 2006). The levels of THC and CBD in cannabis vary depending on the cultivar, and consumers often seek out specific ratios based on their reason for use. According to a recent study examining over 700 people, most people who use cannabis recreationally report using high THC, low CBD cultivars, people who use cannabis medically most often use low THC, high CBD cultivars, and those who use cannabis recreationally *and* medically report using cultivars with a wide range of THC:CBD ratios (Turna et al., 2020). In addition, the vast majority of cannabis users report not knowing what the percentage of THC or CBD is in the products they consume (Turna et al., 2020).

It has been shown that THC, CBD, and the combination of the two phytocannabinoids produce differing effects on the body and brain, likely through unique influences on the endocannabinoid system through different receptors (Zou and Kumar, 2018). THC is a partial agonist at the cannabinoid type 1 receptor (CB1R)—the most common endocannabinoid receptor in the central nervous system (CNS) and most common G-protein coupled receptor in the brain (Thomas et al., 2007; Pertwee, 2008; Laaris et al., 2010; Laprairie et al., 2015). The vast majority of CB1R are found in the presynaptic terminal of gamma-aminobutyric acid (GABA) inhibitory neurons throughout the brain, where their function is to suppress neurotransmitter release (Herkenham et al., 1990; Bonilla-Del Río et al., 2021; Chou et al., 2022). However, CB1R is also present on the presynaptic terminal of glutamatergic neurons (where they also suppress neurotransmitter release), mitochondria, and glial cells, where they function to alter cellular metabolism, cell-cell communication, phagocytosis, and more (Steindel et al., 2013; Busquets-Garcia et al., 2018; Bonilla-Del Río et al., 2021; Murray et al., 2023). In addition, THC also interacts with many other receptors such as G protein-coupled receptor 55 (GPR55: agonist), but the majority of its effect is thought to be driven by CB1R (Lauckner et al., 2008; Stollenwerk et al., 2021). The cannabinoid type 2 receptor is also thought to be present in the brain at much lower levels, mainly expressed on microglia, where THC acts as a partial agonist (Pertwee, 2008; Jordan and Xi, 2019).

Compared to THC, CBD is more enigmatic. While CBD may act as an indirect antagonist or negative allosteric modulator at CB1R, at physiologically relevant levels CBD likely has minimal direct impact on CB1R function (Laprairie et al., 2015; Peng et al., 2022). However, CBD has a higher affinity for many other receptors/proteins, such as GPR55 (antagonist), adenosine A_2A_ receptor (indirect agonist via inhibition of equilibrative nucleoside transporter-1 [ENT-1]), transient receptor potential vanniloids (TRPVs; agonist), peroxisome proliferator-activated receptor gamma (PPARγ; agonist), 5-hydroxytryptamine 1_A_ (5-HT_1A_; agonist–under debate), and fatty-acid amide hydrolase (FAAH; inhibitor), the enzyme responsible for breaking down the endocannabinoid anandamide (AEA) (Carrier et al., 2006; Peng et al., 2022; Puighermanal et al., 2024; Alexander et al., 2025). Given its pleiotropic targets in the CNS, it is difficult to describe through which pathway CBD exerts its proposed effects (Stella, 2023). CBD has also been shown to potentiate, attenuate, or have little effect on the pharmacokinetics, behavioural impairments, or changes to brain communication caused by THC when administered together in mice, rats, and humans (Roser et al., 2009; Bhattacharyya et al., 2010; Klein et al., 2011; Englund et al., 2013; Todd and Arnold, 2016; Murphy et al., 2017; Boggs et al., 2018; Arkell et al., 2019; Freeman et al., 2019; Nidadavolu et al., 2021; Moore et al., 2023; Zamarripa et al., 2023; Ertl et al., 2024; Chester et al., 2026).

Most cellular investigations examine (phyto)cannabinoid effects on neurons, though recent research suggests (phyto)cannabinoids interact with glia, especially those of the oligodendrocyte lineage. Oligodendrocyte progenitor cells (OPCs) have a multiplicity of functions in the brain beyond their canonical role as a precursor pool to replace or contribute to the mature oligodendrocyte population through oligodendrogenesis. Indeed, a large portion of OPCs never differentiate, but instead persist in the adult brain where they have been shown to prune, engulf and digest synapses and portions of neuronal axons, regulate growth and remodeling of axon arbors, contribute to angiogenesis, actively participate in inflammatory processes and antigen presentation, and more (Kirby et al., 2019; Auguste et al., 2022; Buchanan et al., 2022; Xiao et al., 2022; Xiao and Czopka, 2023; Fang et al., 2025).

Oligodendrocyte lineage cells also express genes essential to the endocannabinoid system in mice, including *Cnr1* (gene for CB1R), and the primary enzymes responsible for the production and degradation of endocannabinoid ligands (*Mgll, Dagl*⍺*, Nape-pld,* and especially *Faah*) (**Supplementary Fig. 1**; Zhang et al., 2014; Yasuda et al., 2020). In line with this, numerous studies have now established a direct and cell-autonomous influence of cannabinoids on OPC function and biology in rodents (Ilyasov et al., 2018; Murray et al., 2023). For example, THC (3 mg/kg; intraperitoneal [*i.p*]) promotes CB1R-dependent OPC cell cycle exit, differentiation, and enhanced myelination in young mice (Molina-Holgado et al., 2002; Huerga-Gómez et al., 2021; Manterola et al., 2022; Sánchez-de la Torre et al., 2022). There are conflicting reports on the effect of CBD on the oligodendrocyte lineage. For example, one study found that CBD (100 nM and 1 μM) may be cytotoxic to primary cultured oligodendrocytes from rats via a rise in intracellular Ca^2+^, whereas another study in primary cultured OPCs from rats found that CBD (1 μM) was protective and increased OPC viability by reducing endoplasmic reticulum stress (Mato et al., 2010; Mecha et al., 2012). However, it is unclear if these doses are physiologically relevant for most individuals who consume CBD via common routes of administration (i.e., inhalation and orally). Additionally, the combined effects of THC and CBD on oligodendrocyte lineage cells have yet to be explored.

To this end, we set out to investigate how cells of the oligodendrocyte lineage in the forceps minor respond to acute cannabis administration of different cannabis cultivars through the inhalation of vaporized cannabis flower in adult male and female mice. We reasoned that the forceps minor would be particularly sensitive to the effects of phytocannabinoids due to it being the primary interhemispheric fibre tract connecting the frontal lobes, which are rich in CB1R and particularly impacted by cannabis use (Yanes et al., 2018; Murray et al., 2023; Stoller et al., 2024). Adult mice were exposed to vaporized cannabis flower for 15 minutes and were subsequently euthanized at 30 minutes post-onset in preparation for multi-modal microscopy experiments. Our goal was to quantify potential differences in how OPCs respond to differing ratios of THC:CBD when THC levels are peaking in the brain, while maintaining sex as a factor (Baglot et al., 2021). We hypothesized that the high THC cultivar would have the most pronounced effect on OPC characteristics, followed by smaller effects for balanced, and finally the CBD cultivar.

## Methods

### Animal Procedures

#### Ethics Statement

All animal procedures relating to acute vapor exposure of cannabis flower by young adult (10 weeks old) mice were ethically reviewed and approved by the Animal Care Committee at the University of Guelph (AUP#4194). All procedures strictly followed the recommendations from the Canadian Council on Animal Care.

#### Cannabis Exposure

Adult (10 weeks old) male and female C57BL/6J mice were acutely exposed to one of three different cultivars of dried cannabis flower acquired from the Ontario Cannabis Store or a control vapor air (**Fig. 1A**). The cultivars consisted of high THC (Wedding Mint: 24–30% THC; ∼0% CBD), balanced THC and CBD (Twd.: 6–12% THC; 5–11% CBD), and high CBD (Pure Sun CBD: 12–18% CBD; ∼0% THC) cannabis flower, while the control vapor air consisted of running the vaporizer with no cannabis flower present. Mice were individually placed into 16.4 L self-administration vapor chambers (La Jolla Alcohol Research Inc.) and received a 15-second puff of cannabis flower (0.15 g cannabis flower/puff) or control vapor every 5 minutes for a total of 15 minutes (0.45 g cannabis flower total).

**Figure. 1.**
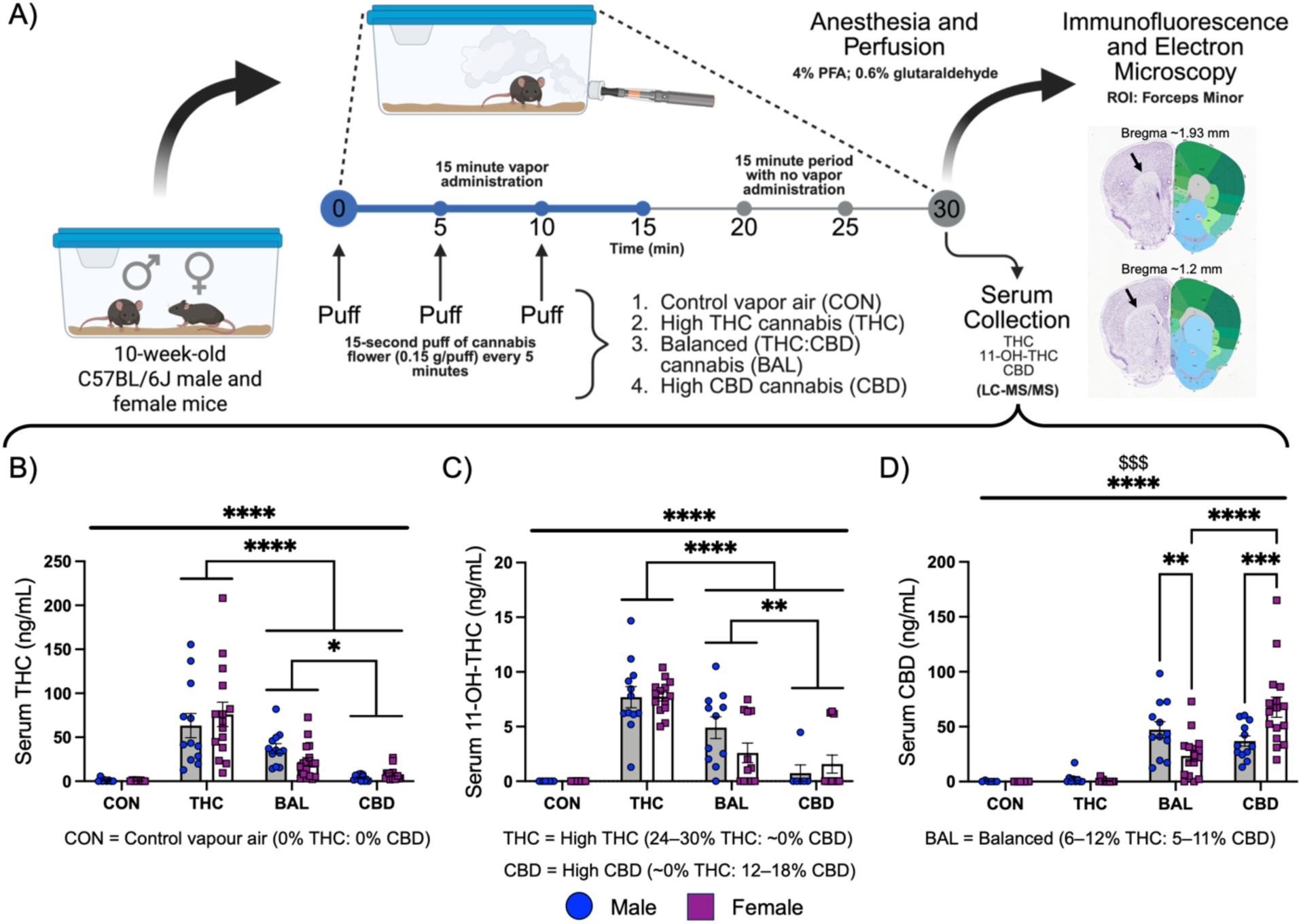
Exposure paradigm and experimental timeline alongside serum levels of THC, 11-OH-THC, and CBD from adult male and female mice acutely exposed to vapor from different cultivars of cannabis and a control vapor air. (A) Mice were exposed to vapor from 0.45 g of cannabis flower over 15 minutes and were subsequently anesthetized and perfused with a fixative solution after 30 minutes post-cannabis onset. Representative images from the Allen Brain Atlas are provided for the beginning and end Bregma range used in this study, and the forceps minor region of interest (ROI) is indicated by black arrows. Cannabis cultivars consisted of high THC, balanced in THC and CBD (BAL), and high CBD. Serum was collected at 30 minutes post cannabis vapor onset via intracardiac puncture, and was analyzed using liquid chromatography-tandem mass spectrometry (LC-MS/MS) (**Supplementary File 1**). For ease of visualization, only select multiple comparisons are shown for serum data: THC *vs.* BAL *vs.* CBD and within groups comparisons where appropriate. We observed significant differences in serum (B) THC, (C) 11-OH-THC, (D) and CBD between the different cultivars and with the control vapor air. All statistical comparisons are available in Supplementary File 2. Serum data (B, C, D) are expressed as group mean ± standard error of the mean. N=6–16 animals/group/sex. Blue circles=male; Purple squares=female. * corresponds to cannabis effect while $ corresponds to an interaction. *p*-value ≤ 0.05*, *p*-value ≤ 0.01**, *p*-value ≤ 0.001***, *p*-value ≤ 0.0001****; *p*-value ≤ 0.05^$^. CON=control; THC=high THC cultivar; BAL=balanced cultivar; CBD=high CBD cultivar. Panel ‘A’ created in Biorender.com.

#### Serum Collection and Cannabinoid Quantification

Immediately preceding the perfusion, a 300 µL blood sample was collected through intracardiac puncture, transferred to Eppendorf tubes, and placed on ice. Blood was centrifuged at 8000 x *g* for 5 min, and serum supernatant was collected and stored at -80°C until analysis. Serum levels of THC, CBD, and 11-hydroxy-THC (11-OH-THC), an active metabolite of THC, were evaluated using liquid chromatography-tandem mass spectrometry (LC-MS/MS) as previously mentioned (Amissah et al., 2024; Kayir et al., 2025). For full LC-MS/MS protocol, see **Supplementary File 1**.

#### Intracardiac Perfusion and Fixation

30 minutes post-onset of cannabis exposure, animals were anesthetized with isoflurane before being intracardially perfused with a peristaltic pump at a speed of 24 mL/min for 10 minutes. Blood was initially flushed with ice-cold phosphate-buffered saline (PBS: 50 mM, pH 7.4) solution, followed by a fixative solution containing 4% paraformaldehyde and 0.6% glutaraldehyde. Brains were dissected and post-fixed in 4% paraformaldehyde for 2 hours at 4°C, followed by PBS washes. Brains were coronally sectioned at a thickness of 50 µm on a vibratome (VT1200S, Leica Biosystems) in ice-cold PBS and subsequently stored at -20°C in cryoprotectant/antifreeze solution (30% (v/v) glycerol: 30% (v/v) ethylene glycol: 40% (v/v) PBS) until further processing.

### Immunofluorescence Microscopy

#### Staining

Brain sections containing the forceps minor (Bregma: ∼1.93–1.2 mm) were selected based on the stereotaxic atlas by Paxinos and Franklin (Paxinos and Franklin, 2012). 3–4 brain sections along the selected Bregma range were selected per animal for n=4–5 animals/group/sex, unless otherwise stated.

For quantification of oligodendrocyte lineage cell density, nearest neighbour distance (NND), spacing index, and OPC morphology, brain sections were first washed 5 x 5 minutes in PBS. Sections then underwent antigen retrieval with sodium citrate buffer (10 mM with 0.5% Tween 80, pH 6.0) at 70°C for 40 minutes in a hot water bath. The well plate was allowed to cool for 15 minutes at room temperature before being washed 5 x 5 minutes with PBS. Samples were quenched with 0.1% sodium borohydride for 30 minutes, followed by 3 x 10 minutes PBS washes. Sections were incubated in blocking buffer containing 10% normal donkey serum, 0.5% fish gelatin, and 0.3% Triton X-100 in PBS for 1 hour at room temperature. Sections were incubated overnight at 4°C in a cocktail of primary antibodies in blocking buffer against platelet-derived growth factor receptor alpha (rabbit anti-PDGFR⍺; 1:500 Abcam Cat#203491), oligodendrocyte transcription factor 2 (goat anti-Olig2; 1:400 R&D Systems Cat#AF2418), and adenomatous polyposis coli (mouse anti-CC1; 1:400 Calbiochem Cat#OP80).

The following day, sections were allowed to reach room temperature before being washed with PBS containing 0.3% Triton X-100 (PBS-T) for 5 x 5 minutes. Sections were incubated in a cocktail of fluorescent secondary antibodies in blocking buffer, including donkey anti-goat Alexa Fluor^®^ 488 (1:300 Invitrogen Cat#A11055), donkey anti-mouse Alexa Fluor^®^ 555 (1:300 Invitrogen Cat#A31570), and donkey anti-rabbit Alexa Fluor^®^ 647 (1:500 Invitrogen Cat#A31573) for 2 hours at room temperature. Following 5 x 5 minutes washes in PBS-T, sections were incubated with 4′,6-diamidino-2-phenylindole (DAPI; 1:20k Invitrogen Cat#D3571) for 5 minutes. Sections were rinsed with phosphate buffer (PB; 10 mM, pH 7.4) 5 x 5 minutes, mounted, and coverslipped with Fluoromount G (Invitrogen Cat#00-4958-02).

A similar protocol with slight modifications was performed for immunofluorescent staining against antigen Kiel 67 (rat anti-Ki67; 1:500 Invitrogen Cat#14-5698-82) alongside PDGFR⍺ (rabbit anti-PDGFR⍺; 1:500 Abcam Cat#203491). Briefly, the differences include antigen retrieval in sodium citrate buffer at 95°C for 30 minutes, washes in PBS-T (0.5%) for all washes following the sodium borohydride incubation, blocking buffer components (5% normal donkey serum, 5% normal goat serum, 0.5% fish gelatin, 0.5% Triton X-100), and primary antibody incubation overnight at 37°C. The donkey anti-rat Alexa Fluor^®^ 488 (1:500; Invitrogen Cat#A21208) secondary antibody was used in addition to donkey anti-rabbit Alexa Fluor^®^ 647 (1:500 Invitrogen Cat#A31573) for 2 hours at room temperature. Brain sections had a more restricted Bregma range for these experiments to limit sampling variability, and were selected between Bregma ∼1.5–1.2 mm.

#### Imaging

Immunofluorescent images containing the forceps minor were acquired using a 20x objective lens (numerical aperture of 0.8) with an Axio Imager M2 epifluorescence microscope with Apotome equipped with an AxioCamMR3 camera (Zeiss, Oberkochen, Germany). For density, NND, and spacing index, a single focal plane was imaged, followed by stitching and background subtraction (radius of 100) performed in Zen 3.1 software (Blue Edition, Zeiss). For quantification of OPC sister cells and proliferative OPCs, a stack of 10–12 µm with an interval of 1 µm was acquired to increase the sampling area. Image stacks then underwent orthogonal projection, stitching, and background subtraction with the same aforementioned settings, using Zen 3.1 software (Blue Edition, Zeiss). For OPC morphology analysis, a 63x objective lens (numerical aperture of 1.4) was used to acquire 30 µm image stacks (0.4 µm step size) with a Zeiss LSM-880 confocal microscope in Airyscan mode (Zeiss, Oberochen, Germany). Image stacks then underwent 3D Airyscan processing using Zen 2.6 software (Blue Edition, Zeiss).

### Electron Microscopy

#### Immunofluorescence for Correlative Light Electron Microscopy (CLEM)

One 50 µm brain section per animal (n=4 animals/group) containing the forceps minor (Bregma: ∼1.93–1.2 mm) was selected based on the stereotaxic atlas by Franklin and Paxinos (Paxinos and Franklin, 2012) and washed for 5 x 5 minutes in PBS. Sections were quenched in 0.05% sodium borohydride in PBS for 30 minutes. After 5 x 5 minutes washes in PBS, sections were incubated in blocking buffer (10% fetal bovine serum, 0.5% fish gelatin, 0.01% Triton X-100) for 1 hour. Sections were directly incubated with rabbit anti-PDGFR⍺ (1:500 Abcam Cat#203491) in blocking buffer overnight at 4°C. The following day, sections were allowed to come to room temperature before 5 x 5 minutes washes in PBS-T (0.01%) and subsequent incubation with donkey anti-rabbit Alexa Fluor^®^ 647 (1:800 Invitrogen Cat#A31573) secondary antibody for 2 hours at room temperature. Sections were washed 5 x 5 minutes with PBS-T and incubated with DAPI (1:20k Invitrogen Cat#D3571) for 5 minutes. Finally, sections were washed for 5 x 5 minutes in PB and mounted with Fluoromount G (Invitrogen Cat#00-4958-02) on non-charged slides.

The forceps minor was imaged in its entirety at 20x (numerical aperture of 0.8) with an Axio Imager M2 epifluorescence microscope with Apotome equipped with an AxioCamMR3 camera (Zeiss, Oberkochen, Germany). Subsequent imaging of specific OPCs (12–15 cells/animal) at 63x (numerical aperture of 1.4) with a Zeiss LSM-880 confocal microscope with Airyscan mode (Zeiss, Oberkochen, Germany) of the top 15 µm (1 µm step size) of the tissue provided more in-depth detail of cell location for future correlative imaging with the scanning electron microscope (SEM). After imaging, coverslips and sections were carefully removed from the slide and sections were re-introduced into PB in a well plate in preparation for heavy metal incubation. Orientation of the sections was carefully recorded to ensure the same surface of the tissue would be cut with the ultramicrotome.

#### Heavy Metal Incubation and Electron Microscopy

After 5 x 5 minutes in PB, sections were incubated in 2% aqueous osmium tetroxide (Electron Microscopy Sciences (EMS) Cat#19170) with 3% potassium ferrocyanide (Millipore Sigma Cat#P9387) in PB for 1 hour at room temperature. Sections were washed in 100% PB, 50% PB:50% Milli-Q water, and finally 100% Milli-Q water for 5 minutes each before a 20-minute incubation in 1% thiocarbohydrazide (TCH: Millipore Sigma Cat#223220) in Milli-Q water. Following 3 x 5 minute washes in Milli-Q water, sections were incubated with 2% osmium tetroxide in Milli-Q water for 30 minutes, washed for 3 x 5 minutes Milli-Q water, and subsequently dehydrated in ascending concentrations of ethanol (2 x 35%, 1 x 50%, 1 x 70%, 1 x 80%, 1 x 90%, 3 x 100% for 5 minutes each), and finally in 3 x 5 minutes propylene oxide (Millipore Sigma Cat#110205). Sections were then placed in Durcupan ACM resin (Millipore Sigma Cat#44611-44614) at room temperature overnight. The following day, resin-embedded sections were delicately placed in a thin layer of resin between two sheets of Aclar^®^ embedding films (EMS Cat#50425-25) and polymerized at 55°C for 72 hours. The region of interest was then micro-dissected and glued to a Durupan resin block and cut into ultrathin sections of 74 nm using a Leica ARTOS 3D ultramicrotome (Leica Biosystems). For CLEM experiments, 5–10 74 nm ultrathin sections every ∼1–2 µm from the top 15 µm of the tissue were serial sectioned and collected to allow for correlation with confocal Z-stack images. Ultrathin sections were collected on silicon nitride chips and placed on specimen mounts for the SEM. Samples were imaged at 5 nm resolution (x, y) using a ZEISS Crossbeam 350 Gemini SEM, operating with an acceleration voltage of 1.5 kV, 1.2 nA current and a working distance of 5 mm.

### Data Analyses

All data analyses were conducted blind to experimental conditions.

#### Density and Distribution

20x images acquired with an epifluorescent microscope were analyzed using QuPath (v0.3.2) and ImageJ (v1.53a, NIH) software. In QuPath, the forceps minor in its entirety—with reference to the Paxinos and Franklin Atlas—was traced using the ‘polygon tool’ and then the ‘points tool’ was used to manually count all mature oligodendrocytes (CC1^+^/Olig2^+^), OPCs (PDGFR⍺^+^/Olig2^+^) and cells of the oligodendrocyte lineage (CC1^−^/PDGFR⍺^−^/Olig2^+^) within the region (Paxinos and Franklin, 2012). Cell density values were calculated by dividing the cell count by the traced area (cells/mm^2^). The ‘points’ annotation, which marks each OPC, was then sent to ImageJ through a QuPath extension, and the distribution of OPCs was quantified using the NND plugin (Integrated Computational Materials Engineering, Mississippi State University). 3–4 sections/animal were used for n=4–5 animals/group/sex.

#### OPC Sister Cells and Proliferative OPCs

20x orthogonally projected images taken with an epifluorescent microscope were used for quantifying the density of OPC (PDGFR⍺^+^/Olig2^+^) sister cells, a proxy measure for OPC proliferation. OPCs were considered sister cells based on their mirrored morphology and close apposition of their soma (Boda et al., 2015; Shin and Kawai, 2021). In addition, to only include recent divisions, OPC sister cells were only counted if their somas were within 10 µm of each other—the distance the majority of sister cells are separated by—which was confirmed using manual measurements in QuPath (Boda et al., 2015; Shin and Kawai, 2021). Density analysis was performed identically as described above. To further analyze OPC proliferation, the proportion of Ki67^+^/PDGFR⍺^+^ to Ki67^−^/PDGFR⍺^+^ OPCs within the forceps minor was quantified in QuPath. In addition, the proportion of OPC sister cells double positive for Ki67 (doublets), singly positive (singlets), or negative was also quantified in QuPath. 3–4 sections/animal were used for n=4 animals/group/sex.

#### OPC Morphology

63x Airyscan image stacks taken with a confocal microscope were analyzed using Imaris software (v10.0.2, Oxford Instruments). OPCs were randomly selected across 3–4 sections per animal for a total of 15 cells/animal for this analysis (n=4 animals/group/sex). We reconstructed cells in an identical manner as performed previously (Murray et al., 2025). Of note, the thinnest filament diameter used in the filaments tool was set to 0.5 µm, and the seed point threshold was manually set so that all visible branches had at least one point at their base and tip. Reconstructed cells were analyzed using the ‘Filament Sholl Analysis 24’ plugin, developed by Ironhorse1618 (2023): https://github.com/Ironhorse1618/Python3.7-Imaris-XTensions with a Sholl radius set at 10 µm. Under the ‘Statistics’ tab in Imaris, the quantified values of each measurement of interest were acquired (**Supplementary File 2**).

Respective morphology indices (i.e., total, soma, branching, and convex hull morphology index) were calculated by first performing a principal component analysis (PCA) with all measures within each ‘index’ using GraphPad Prism (v10.0.3 Graphstats Technologies). Principal components (PCs) were chosen if they had eigenvalues ≥1, and their sum needed to account for at least 75% of the variance within the dataset (**Supplementary File 2**). If 75% variance was not achieved with the PCs with eigenvalues ≥1, an additional PC was included. The proportion of variance of each PC was then normalized to the total variance explained by the selected PCs to get the weighted variance (*w*) explained by each PC. The weighted variance of each PC was then multiplied by their corresponding PC score (*p*) to create a weighted PC score per component. The weighted scores were then summed across all selected PCs to create the final weighted index score (*morphology index*) for each cell.

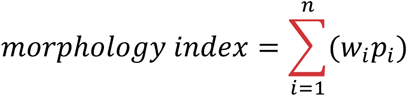

#### Ultrastructural Analyses

10–20 oligodendrocytes/animal were identified and randomly selected for analysis. Oligodendrocytes were identified based on their unique ultrastructural characteristics: square-shaped nucleus, cytoplasm pushed to one pole of the cell, electron-dense cytoplasm, lack of intermediate filaments or glycogen granules (both prominent in astrocytes), as well as short and wide endoplasmic reticulum cisternae (Peters, 1991; Nahirney and Tremblay, 2021). They also often formed ‘trains’ of oligodendrocytes in the white matter (Peters, 1991). Satellite oligodendrocytes on blood vessels were not included in the analysis.

To aid in the identification of OPCs, we utilized CLEM with PDGFR⍺ and DAPI. 12–15 OPCs/animal were randomly selected and imaged with a confocal microscope before tissue preparation for SEM. Using landmarks (i.e., blood vessels) and the distribution of DAPI, we were able to identify many of the exact same PDGFR⍺^+^ OPCs from our confocal images. Using the ultrastructural characteristics of these confirmed OPCs, as well as documented characteristics in the literature (i.e., oblong nucleus, relatively low levels of heterochromatin compared to microglia, relatively electron-lucent cytoplasm compared to microglia, and absence of glycogen granules compared to astrocytes), we then selected further cells to reach a sample size of 11–20 OPCs/animal without the use of CLEM (Mori and Leblond, 1970; Buchanan et al., 2022).

The heterochromatin pattern of each nucleus was segmented using a custom U-Net convolutional neural network implemented in PyTorch (Ronneberger et al., 2015; Paszke et al., 2019). The model was trained on 200 manually traced heterochromatin pattern nuclear images and achieved an average training loss of 0.1838 and a mean Dice coefficient of 0.9471 (8 epochs; batch size = 4; learning rate = 0.001) (**Supplementary Fig. 2**). Heterochromatin pattern was quantified from the predicted binary masks using custom Python scripts. Within the nucleus, heterochromatin area was calculated as the number of black pixels in the binarized mask and converted to µm² based on the pixel size (5 nm/pixel). Chromatin coverage was defined as the percentage of nuclear area occupied by heterochromatin. To assess spatial organization, lacunarity was computed across multiple box sizes using the gliding-box algorithm (Allain and Cloitre, 1991). To summarize the scale dependence of heterochromatin heterogeneity, a linear regression was performed between ln(lacunarity) and ln(box size), and the resulting slope was used for statistical comparison. Fractal dimension was estimated using a box-counting approach across multiple spatial scales by regressing ln(N) against ln(1/box size) (Sarkar and Chaudhuri, 1994; Jelinek et al., 2013). For both analyses, only boxes intersecting the region of interest (ROI) were included. To further quantify changes to chromatin distribution, we subdivided the nucleus into non-overlapping concentric shells (0.5 µm thickness) extending inward from the nuclear boundary in Python, using a similar logic published previously (Cherkezyan et al., 2014). The total fraction of each shell covered by heterochromatin was then quantified using the binary mask. Area under the curve (AUC) for chromatin fraction per shell was calculated in GraphPad Prism.

Mitochondria and autophagosomes within the cell soma and the width of nuclear pores were traced manually in QuPath. Annotations were then sent to ImageJ for calculation of shape descriptors including area, Feret diameter, circularity, roundness, and aspect ratio. Mitochondria density was calculated by dividing the number of mitochondria by the area of cytoplasm in the soma (excluding the nucleus). Similarly, mitochondria area fraction was calculated by dividing the total summed area of all mitochondria by the area of the cytoplasm.

### Statistical Analyses

Approximate data normality was assessed visually using QQ plots. Phytocannabinoid and metabolite serum data were analyzed using a two-way analysis of variance (2-way ANOVA). Select multiple comparisons between sex within each cannabis group and between controls and each cannabis group within each sex were performed if a significant interaction was observed, and were corrected with the Šídák method. If no significant interaction was observed, but a main effect of cannabis type was present, *post-hoc* comparisons were performed between each cannabis group, collapsing across sex, and were corrected for with the Tukey method (Rich-Edwards and Maney, 2023).

All density, distribution, and proportional cell data (i.e., % cell) were analyzed using a 2-way ANOVA, following the same *post-hoc* guidelines as stated above. These tests were performed using animal means calculated from technical replicates (3–4 brain sections/animal, n=3–5 animals/groups/sex) in GraphPad Prism (v10.0.3 Graphstats Technologies). In addition, for these analyses and the following (i.e., morphology data), effect size (mean difference [*Δ*] or Cohen’s d [*d*]) and 95% confidence intervals (CI) were calculated for specific comparisons, including between control and cannabis groups within each sex, and between control male and control female groups.

The proportional differences between the three categories of Ki67^+/–^ OPCs in Figure 3 were analyzed in R (v2024.04.2+764) using a Pearson’s chi-square test with Rao-Scott adjustment (survey package: Lumley, 2024) to account for the nested structure of the data and non-independent data points within animals. Multiple comparisons were corrected for using the Šídák method.

OPC morphology and electron microscopy data were analyzed in R. To more accurately take into account animal variability within groups, we opted to use a nested statistical approach, with animals (15 cells/animal, n=4 animals/group/sex) nested within cannabis exposure and sex, as previously performed (Murray et al., 2025). In addition, for the analysis of mitochondria, all mitochondria within a cell were measured and nested within their respective cells, and cells were nested within animals (1|Animal/Cell) to account for the non-independence of mitochondrial measurements within each cell. We used a linear mixed-effects model where cannabis exposure and sex are fixed factors and animal is a random factor, utilizing random intercepts. For measures where data were bounded between 0 and 1, we used a generalized linear mixed-effects model with a beta family distribution.

We performed likelihood ratio tests to assess for significant interactions or main effects (Winter, 2013). We then computed the estimated marginal means for the combinations between cannabis exposure and sex, followed by custom pairwise contrasts with Šídák correction, where appropriate. The Kenward-Roger degrees-of-freedom method was used to estimate the degrees of freedom. RStudio packages were utilized as previously reported (Murray et al., 2025). In addition, model-based group differences (*Δ*) and their 95% CI were calculated using parametric bootstrapping in R. For Index data, effect sizes are expressed as Cohen’s d with 95% CI.

For all analyses, statistical significance was considered when *p*-value ≤ 0.05, and marginal differences were reported at *p*-value ≤ 0.1 (no *post-hoc* was performed when *p* ≤ 0.1); where * denotes significant findings for cannabis exposure, # denotes significant findings for sex, and $ denotes a significant interaction. \**p* ≤ 0.05, \*\**p* ≤ 0.01, \*\*\**p* ≤ 0.001, \*\*\*\**p* ≤ 0.0001, and #*p* ≤ 0.05, ##*p* ≤ 0.01, ###*p* ≤ 0.001, ####*p* < 0.0001. Cohen’s d was reported at *d* ≥ 0.5. All figures were created in GraphPad Prism and RStudio.

Bar graphs are shown as mean ± standard error of the mean (S.E.M.). Raincloud plots for OPC morphology and mature oligodendrocyte data are expressed as model-based means with their 95% Wald CI, alongside the distribution of raw data and individual animal means. OPC SEM data are expressed as mean ± S.E.M. for each animal, with averaged cell values showing variability within animals.

All statistical analyses and outcomes are available in **Supplementary File 3**.

## Results

### Phytocannabinoid serum levels from different cultivars of cannabis have sex-dependent effects

We first quantified the serum levels of THC, its primary active metabolite 11-OH-THC, and CBD 30 minutes after the onset of cannabis flower vapor inhalation of different cannabis cultivars (**Fig. 1A**; **Supplementary File 3**). For serum levels of THC, we observed a main effect of cannabis type (*F*_(3, 92)_ = 28.87, *p* < 0.0001), but no main effect of sex or significant interaction between cannabis and sex (**Fig. 1B**). *Post-hoc* analysis revealed serum from THC groups contained significantly higher serum THC levels than serum from CON (*p* < 0.0001; *Δ* = 68.84 ng/mL; 95% CI 45.08, 92.6), BAL (*p* < 0.0001; *Δ* = 40 ng/mL; 95% CI 19.7, 60.3), and CBD (*p* < 0.0001; *Δ* = 63.73 ng/mL; 95% CI 43.3, 84.16) groups. In addition, BAL groups had significantly higher serum THC compared to CBD (*p* = 0.01; *Δ* = 23.73 ng/mL; 95% CI 3.576, 43.89) and CON (*p* = 0.01; *Δ* = 28.84 ng/mL; 95% CI 5.318, 52.36) groups.

For serum levels of 11-OH-THC, we observed a main effect of cannabis type (*F*_(3, 79)_ = 43.44, *p* < 0.0001) and no main effect of sex or significant interaction between cannabis type and sex (**Fig. 1C**). In line with the THC data, our *post-hoc* analysis revealed significantly higher serum 11-OH-THC in the THC groups compared to CON (*p* < 0.0001; *Δ* = 7.72 ng/mL; 95% CI 5.78, 9.65), BAL (*p* < 0.0001; *Δ* = 3.97 ng/mL; 95% CI 2.12, 5.81), and CBD (*p* < 0.0001; *Δ* = 6.56 ng/mL; 95% CI 4.51, 8.6) groups. We also observed significantly more serum 11-OH-THC in the BAL groups compared to CBD (*p* = 0.009; *Δ* = 2.59 ng/mL; 95% CI 0.5, 4.69), and CON (*p* < 0.0001; *Δ* = 3.748 ng/mL; 95% CI 1.759, 5.737), groups.

Lastly, for serum levels of CBD, we observed a significant interaction between cannabis type and sex (*F*_(3, 83)_ = 7.951, *p* = 0.0001) and a main effect of cannabis type (*F*_(3, 83)_ = 32.51, *p* < 0.0001), but no main effect of sex (**Fig. 1D**). Our *post-hoc* analysis revealed significantly more serum CBD in the CBD and BAL group compared to both the THC and CON groups for both sexes. The female CBD group had significantly more serum CBD than the male CBD (*p* = 0.0002; *Δ* = 30.97 ng/mL; 95% CI 15.09, 46.86) and the female BAL (*p* < 0.0001; *Δ* = –44.35 ng/mL; 95% CI –63.74, –24.96) groups. In addition, the male BAL group had significantly higher serum CBD than the female BAL group (*p* = 0.004; *Δ* = 23.83 ng/mL; 95% CI 7.942, 39.71), but relatively similar CBD levels compared to the male CBD group (*p* = 0.61; *Δ* = 10.45 ng/mL; 95% CI –11.94, 32.83).

In summary, other than the expected differences between cannabis cultivars, we observed sex-dependent effects for serum levels of CBD. In particular, the male BAL mice had higher serum levels of CBD compared to female BAL mice, and the female CBD mice had significantly higher serum CBD compared to male CBD mice.

### Acute cannabis exposure does not have population-level effects on oligodendrocyte lineage cell density and distribution

To investigate the impact of different cultivars of cannabis on oligodendrocyte lineage cells in the forceps minor of male and female mice 30 minutes post cannabis exposure, we first sought to analyze the density of total oligodendrocyte lineage cells (Olig2^+^), mature oligodendrocytes (CC1^+^/Olig2^+^), and OPCs (PDGFR⍺^+^/Olig2^+^) through immunofluorescent microscopy (**Supplementary Fig. 3A**). We did not observe a significant interaction between cannabis type and sex, or a main effect of cannabis type for total oligodendrocyte lineage cells, mature oligodendrocytes, or OPCs (**Fig. 2Bi–iii**). However, we did observe main effects of sex for total oligodendrocyte lineage cells (*F*_(1,29)_ = 11.29, *p* = 0.002), mature oligodendrocytes (*F*_(1,29)_ = 5.79, *p* = 0.02), and OPCs (*F*_(1,29)_ = 9.54, *p* = 0.004). Males had a higher density of total oligodendrocyte lineage cells compared to females, likely driven by a larger population of mature oligodendrocytes, and females had a higher density of OPCs compared to males. We did not observe any significant interactions or main effects of cannabis or sex for OPC NND or spacing index (**Fig. 2E, F**).

**Figure 2.**
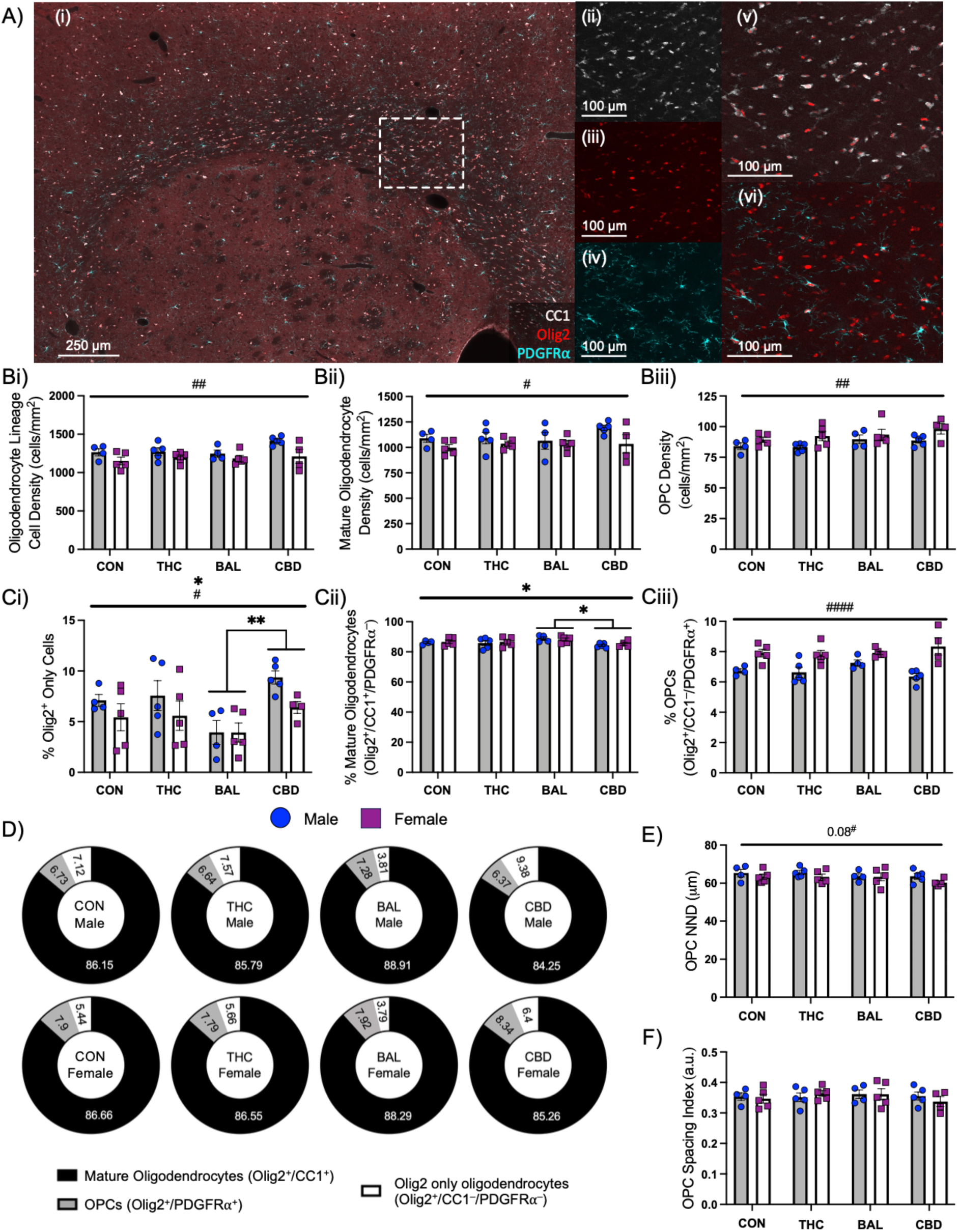
Density and distribution of oligodendrocyte lineage cells do not change in response to acute inhalation of vapor from different cannabis cultivars. (Ai) Representative epifluorescence images of immunofluorescence staining against (Aii) CC1 (white), (Aiii) Olig2 (red), and (Aiv) PDGFR⍺ (cyan), representing (Av) mature oligodendrocytes (Avi) and OPCs. No changes were observed in the density of (Bi) total (Olig2^+^) oligodendrocyte lineage cells, (Bii) Olig2^+^/CC1^+^ mature oligodendrocytes, (Biii) or Olig2^+^/PDGFR⍺^+^ OPCs due to cannabis. Although, there were sex differences, where (Bi) oligodendrocyte lineage cell and (Bii) mature oligodendrocyte density were lower in females compared to males, and (Biii) OPC density was greater in females compared to males. A main effect of cannabis was found for the proportion of (Ci) Olig2^+^/CC1^-^/PDGFR⍺^-^, and (Cii) Olig2^+^/CC1^+^/PDGFR⍺^-^cell populations compared to all oligodendrocyte lineage cells. *Post-hoc* analysis revealed a significant difference between the BAL and CBD groups (*p*-value = 0.01) for Olig2-only cells, and mature oligodendrocytes (*p*-value = 0.02). No main effect of cannabis or interaction was observed for the (Ciii) Olig2^+^/CC1^-^/PDGFR⍺^+^ cell population. (D) Percent breakdown of the entire oligodendrocyte lineage by maturation stage. No change to (E) OPC nearest neighbor distance (NND) or (F) spacing index was observed due to cannabis. Two-way ANOVA with Šídák *post-hoc* where appropriate (B, C, E, F). Data expressed as group mean ± standard error of the mean. N=4–5 animals/group/sex, 3–4 sections/animal. Blue circles=male; Purple squares=female. * corresponds to cannabis effect while # corresponds to sex effect. *p*-value ≤ 0.05*; *p*-value ≤ 0.05^#^, *p*-value ≤ 0.01^##^, *p*-value ≤ 0.0001^####^. CON=control; THC=high THC cultivar; BAL=balanced cultivar; CBD=high CBD cultivar.

Although the density of these different oligodendrocyte populations may not be impacted by cannabis at this time scale, we observed main effects of cannabis type on the proportions of Olig2^+^ only (*F*_(3,29)_ = 4.06, *p* = 0.02), and CC1^+^/Olig2^+^ mature oligodendrocyte (*F*_(3,29)_ = 3.31, *p* = 0.03) populations relative to the total oligodendrocyte lineage pool (all olig2^+^ cells). Our *post-hoc* analysis revealed significant differences between the BAL and CBD groups for the Olig2^+^ only (*p* = 0.01; *Δ* = –3.9%; 95% CI –7.09, –0.8), and mature oligodendrocyte (*p* = 0.02; *Δ* = 3.7%; 95% CI 0.47, 6.9) populations. We also found a main effect of sex for the Olig2^+^ only (*F*_(1,29)_ = 4.24, *p* = 0.05) and OPC (*F*_(1,29)_ = 37.41, *p* < 0.0001) populations, where males showed increased percentage of Olig2^+^ only cells, and females showed increased percentage of PDGFR⍺^+^/Olig2^+^ OPCs.

In summary, cannabis cultivars do not significantly impact oligodendrocyte lineage cell population density or distribution at this time-scale. However, we did observe a significant shift in the proportion of Olig2^+^ only and mature oligodendrocyte populations relative to the whole oligodendrocyte pool in the BAL group compared to CBD group.

### Acute cannabis exposure does not impact the proliferation of OPCs

We next sought to investigate if OPC proliferation may be altered in response to cannabis exposure. To do this, we utilized immunofluorescence microscopy with antibodies against Ki67 and PDGFR⍺ (**Fig. 3A, C**; **Supplementary Fig. 3B**). Initial observations revealed no interaction or main effects of cannabis type or sex for the total percentage of Ki67^+^ OPCs in the forceps minor (**Fig. 3B**).

**Figure 3.**
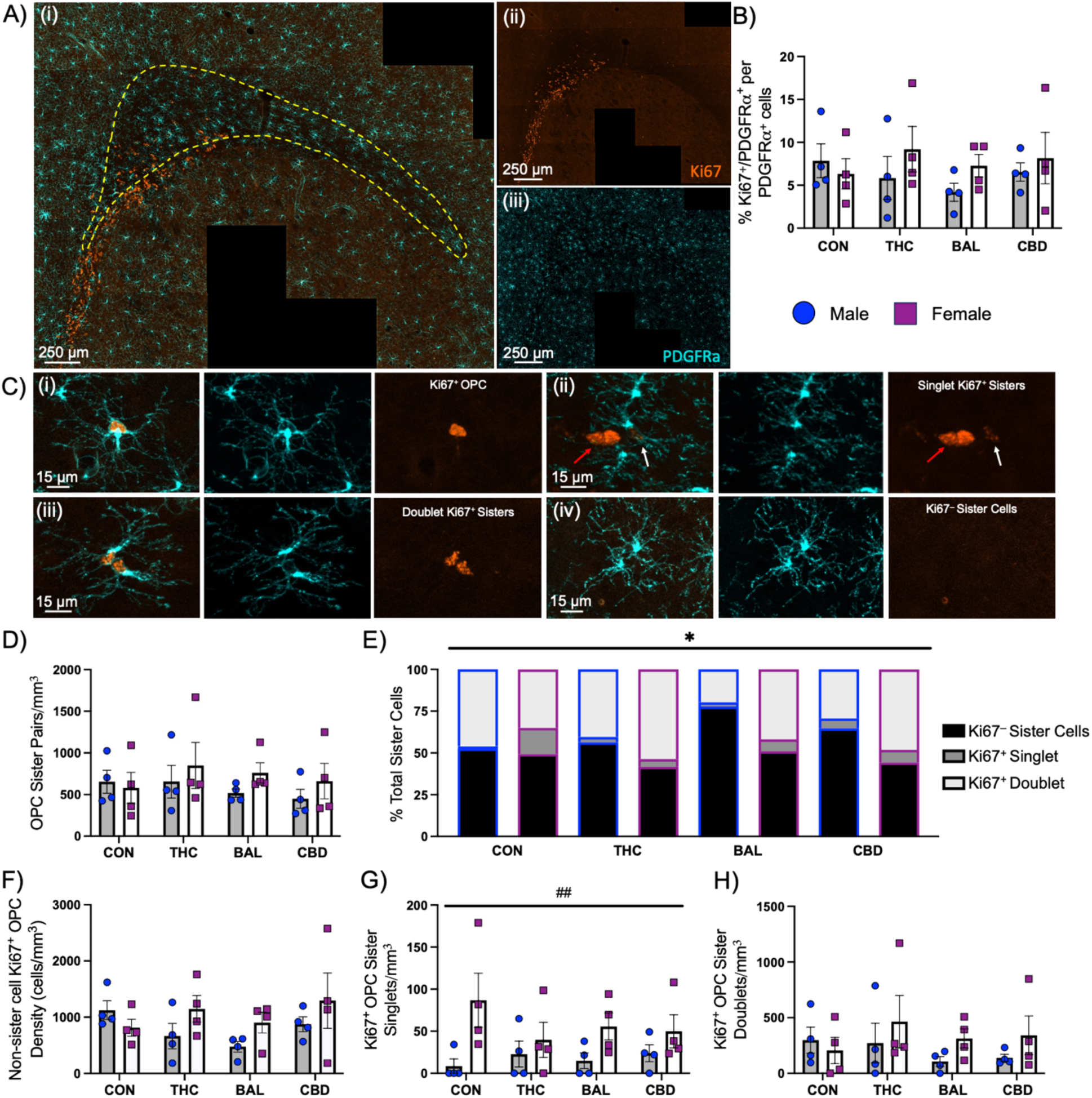
Acute inhalation of vapor from different cannabis cultivars does not alter OPC proliferation or density of OPC sister cells, but may subtly shift OPC sister cell dynamics. (Ai) Representative epifluorescence images of immunofluorescence staining against (Aii) Ki67 (orange) and (Aiii) PDGFR⍺ (cyan). Yellow dotted region outlines the forceps minor, which was traced based off of DAPI staining (not shown). No change in the (B) proportion of Ki67^+^/PDGFR⍺^+^ OPCs compared to the whole PDGFR⍺^+^ population was observed due to cannabis. (C) Representative images of (Ci) Ki67^+^/PDGFR⍺^+^ cells, (Cii) Ki67^+^ singlet sister cells, (Ciii) Ki67^+^ doublet sister cells, and (Civ) Ki67^-^ sister cells. No change to the (D) density of total OPC sister cells, (F) non-sister cell Ki67^+^/PDGFR⍺^+^ cells, (G) Ki67^+^ singlet sister cells, or (H) Ki67^+^ doublet sister cells were observed due to cannabis. (E) A main effect of cannabis (*p*-value = 0.05) was observed for the proportion of OPC sister cells, however, no significant *post-hoc* comparisons were observed. Two-way ANOVA with data expressed as group mean ± standard error of the mean (B, D, F–H). Pearson’s chi-square test with Rao-Scott adjustment with Šídák *post-hoc* (E). N=4 animals/group/sex, 3 sections/animal. Blue circles=male; Purple squares=female. Red arrows in Ciii point to unknown Ki67^+^ cells, whereas white arrows point to Ki67^+^ singlet OPC. * corresponds to effect of cannabis, while # corresponds to sex effect. *p*-value ≤ 0.05*; *p*-value ≤ 0.01^##^. CON=control; THC=high THC cultivar; BAL=balanced cultivar; CBD=high CBD cultivar.

Ki67 is expressed in multiple stages of the cell cycle, especially in G2 and mitosis. Thus, we next aimed to isolate OPCs that have just recently divided from those that may still be in earlier stages of the cell cycle. To do this, we quantified the density of OPC sister cells that were within 10 µm of each other and separated them from Ki67^+^ OPCs that were not part of an adjoining pair. No interaction or main effects of cannabis type or sex were observed for the density of Ki67^+^ OPCs that were not part of an OPC sister pair (**Fig. 3Ci, F**). Similarly, we did not observe a significant interaction or main effects of cannabis type or sex for the density of total pairs of OPC sister cells, irrespective of Ki67 labeling (**Fig. 3D**).

However, recent research has brought to light the asymmetrical nature of OPC division, including with respect to Ki67 expression (Boda et al., 2015; Shin and Kawai, 2021). Therefore, we next aimed to break down our OPC sister cell data into three categories: Ki67^−^ sister cells, Ki67^+^ singlet sister cells (1/2 OPCs Ki67^+^), and Ki67^+^ doublet sister cells (2/2 OPCs Ki67^+^) (**Fig. 3Cii–iv).** We observed a significant overall association between Ki67 category and experimental group (*F*_(6.07,188.17)_ = 2.16; *p* = 0.05; **Fig. 3E**), though we could not identify any group differences with *post-hoc* comparisons. However, it is worth noting that the male BAL group had 26.3% fewer Ki67^+^ doublets compared to CON males, alongside 26.15% more Ki67^−^ sister cells.

We did not observe a significant interaction in terms of the density of Ki67^−^ sister cells, Ki67^+^ singlets, or doublets, or any main effects of cannabis type (**Fig. 3G, H**; **Supplementary File 2**). There was a main effect of sex for Ki67^+^ singlets (*F*_(1,24)_ = 10.01, *p* = 0.004), with female mice having a higher density compared to males (**Fig. 3G**). We also observed a marginal main effect of sex for Ki67^−^ sister cells (*F*_(1,24)_ = 3.894, *p* = 0.06). We did not observe a main effect of sex for Ki67^+^ doublets (**Fig. 3H**).

Together, these data indicate that cannabis exposure does not significantly impact Ki67^+^ OPCs, sister cells, or the proportion of Ki67^+^ OPCs within the greater OPC population at this time scale compared to controls. A subtle effect may be occurring between cannabis exposure and the expression of Ki67 in OPC sister cells.

### Balanced cannabis alters the morphology of OPC soma and nucleus, while CBD-dominant cannabis may influence OPC branching

To identify if different cultivars of cannabis impact OPC morphology, we reconstructed OPCs in 3D using confocal microscopy Z-stacks (**Fig. 4A**). We did not observe a significant interaction or main effects for soma volume, surface area, sphericity, prolate ellipticity, or nucleus:soma volume ratio (**Fig. 4B, C**; **Supplementary Fig. 4B**). However, a main effect of cannabis type was observed for soma oblate ellipticity (χ^2^(3) = 9.66, *p* = 0.02), with a significant *post-hoc* comparison found between THC and CBD (*p* = 0.02; *Δ* = 0.046 a.u; 95% CI 0.012, 0.077) groups. In addition, there was a marginal main effect of cannabis type for soma volume (χ^2^(3) = 6.42, *p* = 0.09) and prolate ellipticity (χ^2^(3) = 6.46, *p* = 0.09).

**Figure 4.**
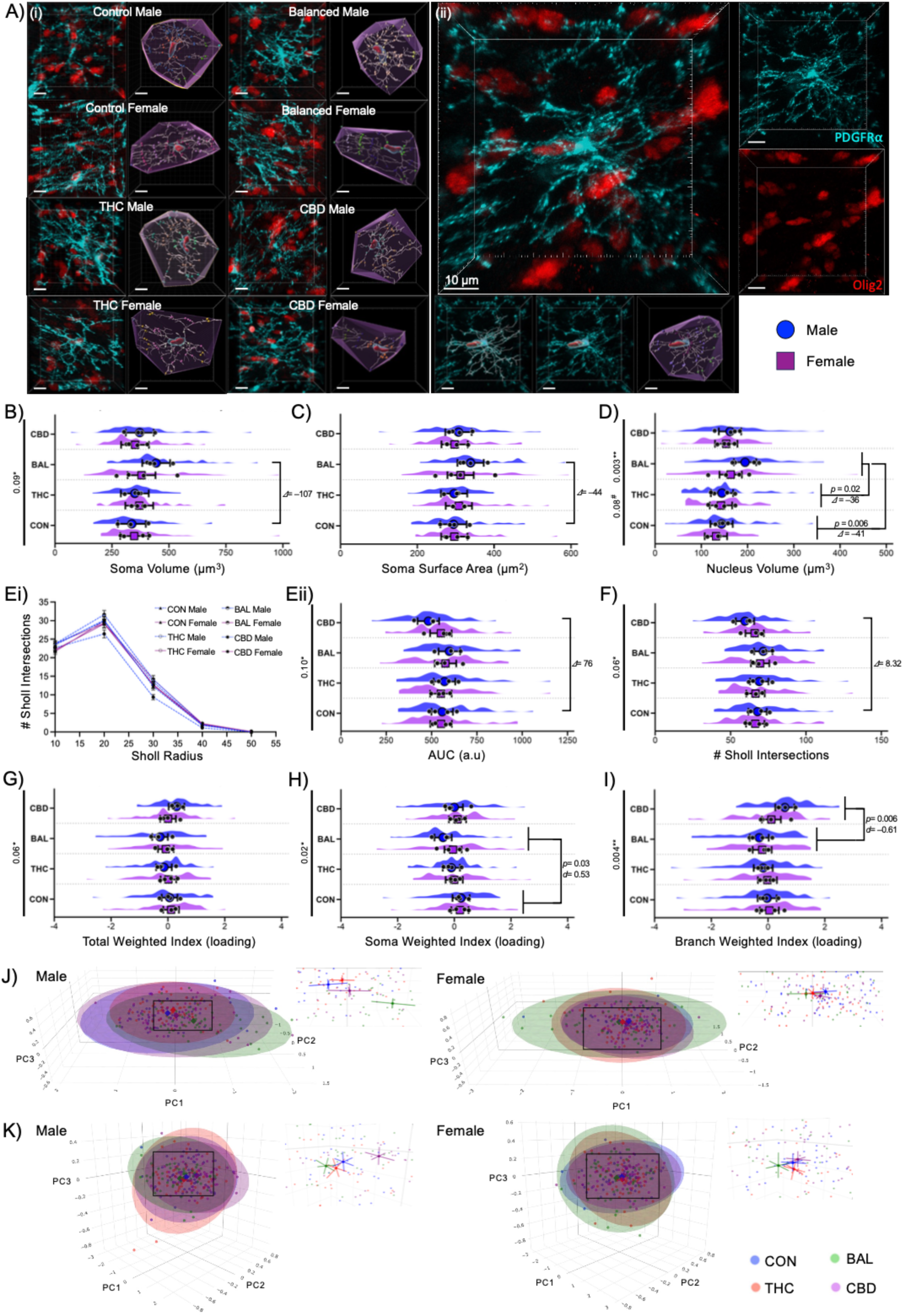
Acute inhalation of vapor from the balanced cannabis cultivar alters OPC morphology in male mice. (Ai) Representative confocal images and 3D reconstruction of nucleus, soma, branches, Sholl intersections and convex hull of OPCs labeled with PDGFR⍺ (cyan) and Olig2 (red) from each experimental group. (Aii) Enlarged representative image of a reconstructed OPC from the balanced male group with separated channels. (B) A potential main effect of cannabis (*p*-value = 0.09) was observed for soma volume, with a large volumetric difference between the male balanced cannabis group compared to control males (*Δ* = –106.78 µm^3^; 95% CI –145.96, –67.99). We observed a (D) main effect of cannabis (*p*-value = 0.003) for nucleus volume. (D) *Post-hoc* analysis revealed a significant increase in nucleus volume (*p*-value = 0.006) in the balanced cannabis group compared to controls (*Δ*= 41.41 µm^3^; 95% CI 20.03, 62.46). No change in (C) soma surface area, (Ei, ii) area under the curve for Sholl analysis, or (F) number of Sholl intersections was observed due to cannabis. We did not observe changes in the (G) total morphological index (including measures from soma, branches and convex hull) or (I) the branching morphological index. (H) We did observe a significant main effect of cannabis (*p*-value = 0.02) for the soma morphological index, alongside significant *post-hoc* comparison between the balanced and control groups. Weighted principal component scores per sex are shown for PC1, PC2 and PC3, which were used to calculate the (J) soma morphological index and (K) branching morphological index. Linear-mixed effects model with data expressed as group model-based mean (blue circle or purple square) ± 95% confidence intervals, alongside raw animal means (black circles) and the spread of raw data from each analyzed OPC (B–D, Eii–I). Šídák *post-hoc* applied to correct for multiple comparisons, where appropriate. N=4 animals/group/sex, 15 cells/animal. Blue circles=male; Purple squares=female. * corresponds to effect of cannabis, while # corresponds to effect of sex. *p*-values shown alongside differences in group means (*Δ*) and Cohen’s D (*d*) for comparisons of interest. CON=control; THC=high THC cultivar; BAL=balanced cultivar; CBD=high CBD cultivar. AUC=area under curve. All scale bars correspond to 10 µm.

We observed a main effect of cannabis type for nucleus volume (χ^2^(3) = 14.09, *p* = 0.003), and a marginal main effect of sex (χ^2^(1) = 2.97, *p*=0.08), but no interaction (**Fig. 4D**). Our *post-hoc* analysis revealed a significant difference between CON and BAL (*p* = 0.006; *Δ* = –41.41 µm^3^; 95% CI –62.46, –20.03) groups, and between THC and BAL (*p* = 0.02; *Δ* = –35.57 µm^3^; 95% CI –55.91, –14.25) groups, with BAL groups showing increased nucleus volume.

We next investigated any possible changes to OPC branch spread through measurements related to convex hull. No significant interactions or main effects were observed for any measurement relating to convex hull, except for those related to sphericity (cannabis type main effect: χ^2^(3) = 9.83, *p* = 0.02; sex main effect: χ^2^(1) = 6.51, *p* = 0.01), where a significant *post-hoc* comparison was found between THC and CBD (*p* = 0.03; *Δ* = –0.014 a.u; 95% CI –0.023, –0.004) groups (**Supplementary Fig. 4A**). In addition, a marginal interaction was identified for prolate ellipticity (χ^2^(1) = 6.9, *p* = 0.08).

To quantify possible changes to OPC branching, we conducted 3D Sholl analysis and plotted a linear regression between the number of Sholl intersections and the corresponding radius. We calculated the AUC and performed a nested statistical analysis. No significant interaction or main effect of sex was found, but we did observe a marginal main effect of cannabis type (χ^2^(3) = 6.27, *p* = 0.10; **Fig. 4Eii**). Similarly, we did not observe any robust effect in any of the other parameters for OPC branching (**Supplementary Fig. 4C**), but we did observe a marginal main effect of cannabis type for the total number of Sholl intersections (χ^2^(3) = 7.39, *p* = 0.07; **Fig. 4F**), Sholl intersections at the 30 µm radius (χ^2^(3) = 6.28, *p* = 0.1), and total filament length (χ^2^(3) = 6.69, *p* = 0.08).

A total weighted morphological index based on individual OPC morphology measures could capture large-scale impacts of cannabis treatment and/or sex. We observed a marginal main effect of cannabis type in the total weighted morphological index (χ^2^(3) = 7.39, *p* = 0.06) (**Fig. 4G).** No significant interaction or main effect of sex was observed.

We further grouped related morphological measures and created an index for each cellular ‘compartment’: soma, convex hull, and branching (**Supplementary File 2**). We observed a main effect of cannabis type on the soma-weighted morphological index (χ^2^(3) = 9.64, *p* = 0.02; **Fig. 4H**). Our *post-hoc* analysis revealed a significant difference between CON and BAL (*p* = 0.03; *d* = 0.53; 95% CI 0.21, 0.85) groups. This shift is likely driven by PC1, which accounts for 44.1% of the variance among the 3 selected PCs, and is strongly influenced by soma volume, surface area, and nuclear volume (**Fig. 4J**, **Supplementary File 2**).

This finding prompted us to go back and take a closer look at the comparison between the CON and BAL groups for soma volume and surface area. For soma volume—the main contributing variable to PC1—although we did not identify a significant interaction or main effect of cannabis type, we observed a small model-based difference in group means between the CON and BAL female groups (*Δ* = –32.32 µm^3^; 95% CI –72.15, 8.93), but a large difference between CON and BAL male groups (*Δ* = –106.78 µm^3^; 95% CI –145.96, –67.99), indicating that the BAL male group may be driving the majority of the effect (**Fig. 4B**). This finding is further bolstered by the fact that the ratio between soma and nucleus volume showed no significant differences (**Supplementary Fig. 4Bi**). Since nucleus size was significantly larger in the BAL group, we would also expect the nucleus:soma ratio to be impacted; however, since the ratio indicates no changes, it suggests that soma volume may be increasing alongside nucleus volume in the BAL group. Indeed, for nucleus volume we observed the largest effect between the CON and BAL male groups (*Δ* = –50.86 µm^3^; 95% CI –70.18, –31.73), and a moderate effect between the CON and BAL female groups (*Δ* = –31.97 µm^3^; 95% CI –51.61, –11.62), showing similar direction of effects to soma volume. In addition, the second highest contributing variable to PC1—soma area—shows a moderate model-based difference in group means (*Δ* = –44.42 µm^2^; 95% CI –67.52, –21.54) between CON and BAL male groups, whereas a smaller effect is observed between CON and BAL female groups (*Δ* = –16.28 µm^2^; 95% CI –39.77, 8.04) (**Fig. 4C**).

We observed a main effect of cannabis type (χ^2^(3) = 13.21, *p* = 0.004) on the branching weighted morphological index (**Fig. 4I**). Our *post-hoc* analysis revealed a significant difference between BAL and CBD (*p* = 0.006; *d* = –0.61; 95% CI –0.92, –0.31) groups. The loadings in our PCA for this index indicated that AUC and total number of Sholl intersections were strongly influencing PC1, which accounted for 44.9% of the variance among the 5 selected PCs (**Fig. 4K**, **Supplementary File 2**). We went back to the nested statistical tests for these measures and took note of the model-based differences in group means. For AUC, we observed a *Δ* = 76.33 a.u (95% CI 23.21, 125.84), and a *Δ* = 8.32 Sholl intersections (95% CI 2.62, 13.62) for total number of Sholl intersections between the CON and CBD male groups (**Fig. 4Eii, F**). However, the model-based group differences for CON *vs* CBD female groups for AUC (*Δ* = 1.17 a.u; 95% CI –52.18, 54.32) and Sholl intersections (*Δ* = 0.4 Sholl intersections; 95% CI –5.32, 6.1) indicate a very small effect.

We did not observe significant effects in our convex hull weighted morphological index (**Supplementary Fig. 4Aiv**).

In summary, we observed significant differences in OPC soma morphology in the BAL cannabis exposed mice, and modest changes to OPC branching due to CBD cannabis. In addition, the effect is particularly pronounced in male mice, which are likely the primary drivers of the main effect of BAL cannabis.

### Balanced cannabis alters OPC mitochondrial density and size at an ultrastructural level

The immunofluorescence data indicated that the OPC soma from animals exposed to the BAL cannabis cultivar is impacted to a greater extent than by other cultivars, particularly in males. To this end, we performed CLEM to confidently identify PDGFR⍺^+^ OPCs, in conjunction with documented ultrastructural characteristics of OPCs, using SEM (**Fig. 5Ai–iv**). Since we only observed robust changes to OPCs between CON and BAL male groups in our immunofluorescence data, we decided to specifically focus on this comparison for the ultrastructural analysis of OPCs.

**Figure 5.**
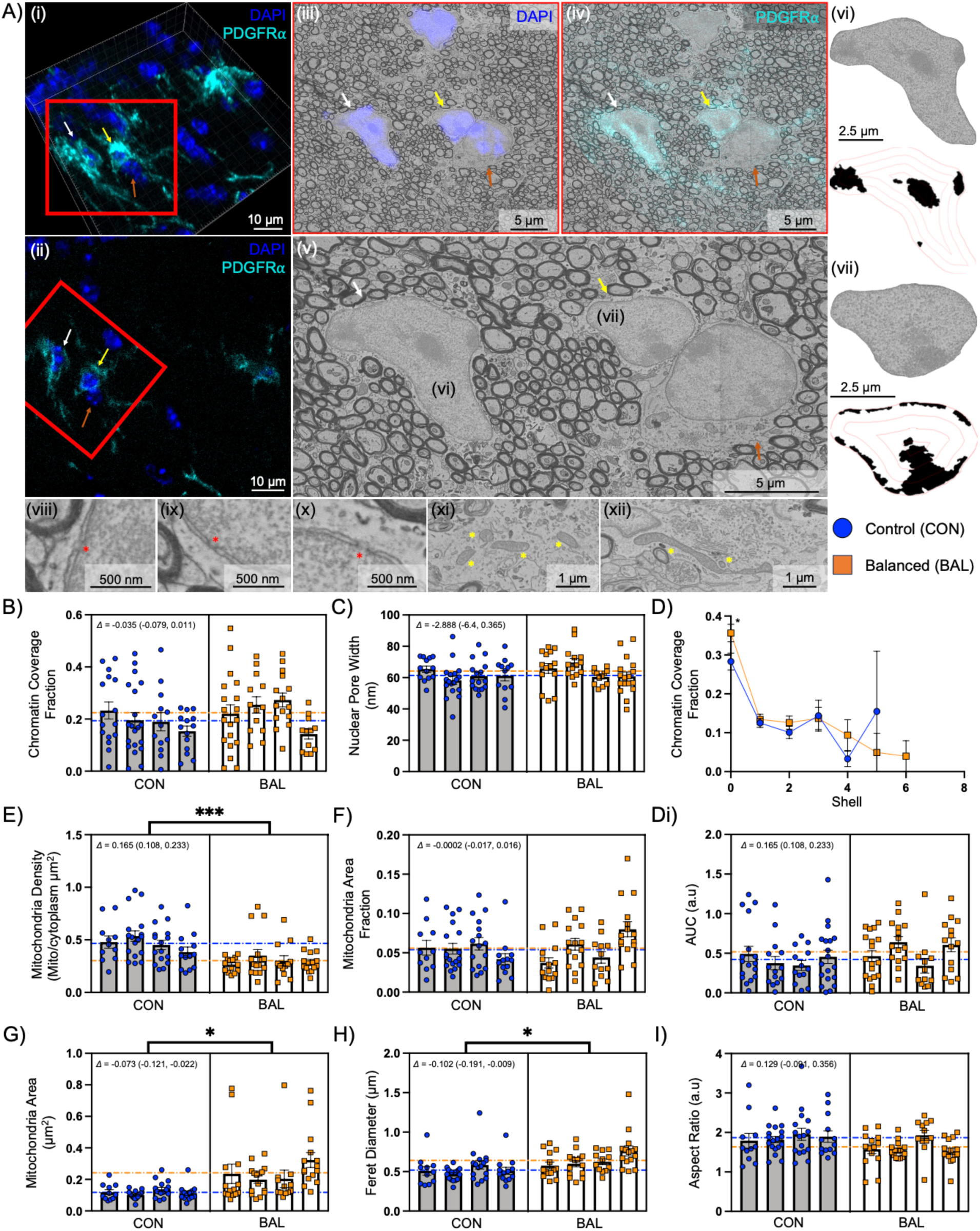
Acute inhalation of vapor from the balanced cannabis cultivar alters mitochondria in OPCs at an ultrastructural level. (Ai) Representative 3D confocal image of immunofluorescence against PDGFR⍺^+^ (cyan) and DAPI (blue), and a (Aii) 2D single plane from the same confocal Z-stack. The red square roughly corresponds to the area visualized in the scanning electron microscope (SEM) images shown in Aiii–v of the same OPCs, using correlative light and electron microscopy (CLEM). (Aiii, iv) DAPI and PDGFR⍺ immunofluorescence are overlaid onto SEM images to show their correlation. (Avi, vii) Representative images of OPC nuclei and their segmentation are shown for the same OPCs seen in images Ai–v. Representative images of (Aviii–x) OPC nuclear pores (marked by red asterisks) and (Axi–xii) mitochondria (marked by yellow asterisks) are shown. (B–Di) We observed no changes to nuclear pore width, or overall heterochromatin coverage due to cannabis. (D) However, we did observe a significant difference between the amount of chromatin coverage in the outermost shell of the nucleus, with the balanced cannabis group having more heterochromatin (*p*-value = 0.05; *Δ* = –0.07 a.u; 95% CI –0.1491, 0.0009). We also observed a significant decrease (*p*-value = 0.0008; *Δ* = –0.165 mito/soma µm^2^; 95% CI –0.233, –0.108) in (E) mitochondria density in the male balanced cannabis group compared to male controls, and significant increases in (G) mitochondria area (*p*-value = 0.02; *Δ* = 0.073 µm^2^; 95% CI 0.022, 0.121) and (H) Feret diameter (*p*-value = 0.05; *Δ* = 0.102 µm; 95% CI 0.009, 0.191). We did not observe any significant differences in (F) mitochondria area fraction or (I) aspect ratio. Linear-mixed effects model with data expressed as animal mean ± standard error of the mean. For mitochondria data, mitochondria are nested within cell, and cell nested within group. Each bar represents one animal, and data points represent the average value for each cell analyzed. Dotted lines correspond to raw group averages. N=4 animals/group/sex, 11–20 cells/animal. Blue circles=control male; Orange squares=balanced cannabis males. (A) White, yellow, and orange arrows point to the same cells in Ai–v. * corresponds to the effect of cannabis. *p*-value ≤ 0.05*; *p*-value ≤ 0.001***. Differences between group means (*Δ*) shown alongside 95% confidence intervals. CON=control; BAL=balanced cultivar. AUC=area under the curve.

We first sought out to identify whether any macro-level changes to the heterochromatin pattern of OPCs were present (**Fig. 5Avi, vii**), which would indicate changes to cell state and/or lineage progression. However, we did not observe any significant effect of BAL cannabis to overall chromatin coverage or distribution, as measured by AUC of heterochromatin coverage per shell (**Fig. 5B, D, Di**), lacunarity (**Supplementary Fig. 5C**), lacunarity slope (**Supplementary Fig. 5D**), and fractal dimension (**Supplementary Fig. 5E**). We did observe a small effect (*p* = 0.05; *Δ* = – 0.0741 a.u; 95% CI –0.149, 0.0009) for chromatin coverage in the outermost shell of the nucleus, which typically has a high percentage of heterochromatin in OPCs, where OPCs from the BAL group had a higher proportion of heterochromatin coverage compared to CON (**Fig. 5D).** We did not observe any significant effect of BAL cannabis on nuclear pore width (**Fig. 5Aviii–x, C).**

We also quantified these measures for mature oligodendrocytes across all groups (**Supplementary Fig. 6**). We observed a main effect of sex for chromatin coverage (**Supplementary Fig. 6F**), with female mice tending to have lower heterochromatin coverage compared to males (χ^2^(1) = 4.44, *p* = 0.04). No other significant effect was observed for mature oligodendrocytes.

Since (phyto)cannabinoids and acute cannabis exposure are known to impact mitochondrial function, we analyzed OPC mitochondria to gain tangential insight into OPC metabolism (Powlowski et al., 2024). We observed a significant difference in OPC mitochondria density, with reductions observed in the BAL group (χ^2^(1) = 11.20, *p* = 0.0008; *Δ* = 0.165 mitochondria/soma µm^2^; 95% CI 0.108, 0.233; **Fig. 5E**). We also observed an increase in mitochondria area (χ^2^(1) = 5.27, *p* = 0.02; *Δ* = –0.073 µm^2^; 95% CI –0.121, –0.022) and mitochondria Feret diameter (χ^2^(1) = 3.81, *p* = 0.05; *Δ* = –0.102 µm; 95% CI –0.191, –0.009) (**Fig. 5G, H**). However, we did not see any significant effect on mitochondrial shape, including roundness, circularity, or aspect ratio (**Supplementary Fig. 5A, B**; **Fig. 5I**), and no change in the mitochondrial area fraction (**Fig. 5F**).

In mature oligodendrocytes, we observed a marginal interaction between cannabis type and sex (χ^2^(3) = 6.6803, *p* = 0.08) for mitochondrial area, and a marginal main effect of cannabis type (χ^2^(1) = 6.936, *p* = 0.07) for mitochondrial circularity (**Supplementary Fig. 6K, N**). No other significant effect was observed for mature oligodendrocyte mitochondria.

Lastly, to help relate our results to the maturation stages of oligodendrocyte lineage cells, we compared heterochromatin coverage and distribution, as well as mitochondrial prevalence and shape between OPCs and mature oligodendrocytes from CON males. We observed significant increases in heterochromatin coverage (χ^2^(1) = 21.22, *p* < 0.0001), AUC (χ^2^(1) = 14.66, *p* = 0.0001), lacunarity (χ^2^(1) = 8.67, *p* = 0.003), lacunarity slope (χ^2^(1) = 9.82, *p* = 0.002), and fractal dimension (χ^2^(1) = 20.38, *p* < 0.0001) in mature oligodendrocytes compared to OPCs (**Supplementary Fig. 7A–F**). In keeping with previous research, we observed significant increases in mitochondrial density and mitochondrial area fraction in mature oligodendrocytes compared to OPCs (χ^2^(1) = 3.40, *p* = 0.05; χ^2^(1) = 5.20, *p* = 0.02, respectively; **Supplementary 7G, H)** (Bame and Hill, 2024). These changes were observed alongside significant increases in mitochondrial circularity (χ^2^(1) = 7.87, *p* = 0.005) and roundness (χ^2^(1) = 6.86, *p* = 0.009), and a decrease in aspect ratio (χ^2^(1) = 5.72, *p* = 0.02) in mature oligodendrocytes, indicating more circular and punctate mitochondria (**Supplementary Fig. 7K–M**).

In summary, we observed significant ultrastructural changes to OPCs in the BAL cannabis group compared to CON males. We report slight changes to heterochromatin distribution at the border of the nucleus, as well as a significant reduction in mitochondrial density and an increase in mitochondrial area. In addition, significant differences in heterochromatin coverage and mitochondrial dynamics were measured between OPCs and mature oligodendrocytes.

## Discussion

This study set out to understand the acute impact of vapor inhalation from different cultivars of cannabis on oligodendrocyte lineage cells in the forceps minor of adult male and female mice. We were particularly interested in the acute effects of cannabis 30 minutes post-onset, when THC levels are peaking in the brain, as THC has been shown to alter OPC function over longer time-scales, but have not yet been examined acutely. We observed significant cultivar-dependent effects on OPC morphology and ultrastructure, with BAL cannabis having a particularly pronounced effect in male mice, contrary to our original hypothesis. The changes to OPC morphology (i.e., increased nucleus and soma volume) and ultrastructure (i.e., reduced density and increased size of soma mitochondria) in the male BAL cannabis group that we observed are reminiscent of the early changes seen during differentiation, and we hypothesize that BAL cannabis may be initiating this process within 30 minutes.

In support of this hypothesis, it has been previously reported in mice that OPC soma size transiently increases during differentiation, alongside a migration of mitochondria out of the soma towards the processes, before returning to the soma as they mature (Stevens et al., 2002; Xu et al., 2021; Chapman et al., 2023, 2024; Bame and Hill, 2024). However, a main question we had was whether vaporized cannabis might be able to cause these changes to OPCs within the time scale of this study, and whether or not differentiation could be initiated this rapidly.

Firstly, although previous studies have shown that THC levels peak in the brain 30 minutes post-inhalation in rodents, there are detectable levels of THC in blood within seconds, and are presumably detectable in the brain within minutes in rodents and humans (Sharma et al., 2012; Baglot et al., 2021). Acute cannabis inhalation containing THC resulted in a reduction in mitochondrial DNA copy number in blood within 15 minutes in humans (Powlowski et al., 2024). At a cellular level, activation and downregulation of plasma-membrane-bound and mitochondrial CB1R have been observed 30 minutes post injection of THC (5 mg/kg, *i.p.*) in neurons and astrocytes of adult mice (Bonilla-Del Río et al., 2021). However, whether or not OPC differentiation can be initiated this rapidly has yet to be determined. One study found that motor skill learning resulted in a significant increase in newly-formed oligodendrocytes (Enpp6^+^) after just 2.5 hours in the subcortical white matter of adult mice, with an increase in cell density by ∼26% (Xiao et al., 2016). The authors suggest the rapidity of the response is likely underestimated, since Enpp6 is not expressed at the earliest stages of differentiation, and not until after PDGFR⍺ is downregulated (Xiao et al., 2016). The changes we observed due to BAL cannabis could represent the earliest changes during differentiation, before PDGFR⍺ expression is lost. Additionally, the OPCs that are most primed for differentiation do not necessarily express PDGFR⍺, but instead are characterized by GPR17; termed committed OPCs (COPs; Fang et al., 2023). We observed a reduction in the proportion of Olig2^+^ only OPCs relative to the entire oligodendrocyte lineage cell pool in the BAL group compared to the CBD group, alongside increases in mature oligodendrocytes. This change could be due to differentiation of COPs into mature oligodendrocytes.

Actomyosin tension in the cytoskeleton could underlie rapid changes to nucleus and soma volume in OPCs (Lourenço and Grãos, 2016). Actomyosin tension is driven by the contraction or relaxation of F-actin polymers in the cytoskeleton—which by extension influences soma size—and is dependent on the activity of non-muscle myosin II (Lourenço and Grãos, 2016). Importantly, the nucleus is directly linked to this system by the linkers of the nucleoskeleton to the cytoskeleton (LINC)-complex, which physically connects the nuclear lamina to the cytoskeleton and maintains the nucleus:soma volume ratio (Méjat and Misteli, 2010; Wu et al., 2022). Non-muscle myosin II activation and subsequent restriction of OPC cytoskeletal reorganization and differentiation is partly regulated by activation of RhoA/Rho-associated protein kinase (ROCK) signaling (Lourenço and Grãos, 2016). Relevant to this study, Sánchez-de la Torre *et al*. (2022) show that THC induces CB1R-dependent OPC differentiation and oligodendrocyte maturation through a reduction of RhoA/ROCK in mice (Sánchez-de la Torre et al., 2022). RhoA and ROCK reductions occur in just 6 hours, but no earlier timepoint was investigated. Another study in cultured rat hippocampal neurons and Neuro2A cultured neurons observed rapid retraction of F-actin-rich growth cones, neurite retraction, and cell body rounding with bath application of CB1R agonists within minutes, and this effect was CB1R-dependent (Roland et al., 2014). Although this response is likely different in the oligodendrocyte lineage, it is plausible that changes to OPC morphology could be driven by CB1R-mediated changes to RhoA/ROCK signaling and a reduction in actomyosin tension within the timeframe of this study.

We did not observe any strong effect of cannabis on the nuclear heterochromatin distribution, such as that which might be expected during OPC differentiation. However, our investigations were limited to a single focal plane and to macro-scale alterations in heterochromatin pattern, which may not be sensitive enough to identify the fine-scale changes 30 minutes after cannabis exposure. Another limitation of this analysis is that OPCs are highly heterogeneous and include cell states with many different functions strewn throughout the cell cycle (Spitzer et al., 2019; Beiter et al., 2022). This likely contributes to the variability we see in the heterochromatin coverage data, where some OPCs have almost no visible heterochromatin, while others exceed 50% coverage. In contrast, heterochromatin distribution in mature oligodendrocytes was more consistent between cells (**Supplementary Fig. 6F**; **Supplementary Fig. 7A**). Future studies will aim to disentangle the different OPC cell states characterized by markers such as GPR17, as well as maturation stages (i.e., PDGFR⍺ *vs* BCAS1 *vs* CAII), using CLEM (Beiter et al., 2022).

In terms of mitochondria, extensive mitochondrial motility has been shown using *in vivo* imaging in OPCs within 30 minutes in mice, and even noticeable displacement within minutes (Bame and Hill, 2024). As discussed, cannabinoids have rapid and variable effects on mitochondria, which may contribute to cannabinoid-mediated CB1R density, energy metabolism, and social behaviour in mice (Bénard et al., 2012; Jimenez-Blasco et al., 2020; Malheiro et al., 2023). Whether or not OPCs have CB1R on mitochondria has yet to be definitively shown, but their presence on mitochondria in neurons and astrocytes provides promising evidence. In addition, a wide range of effects of CBD on mitochondrial function have been reported (such as PPARγ-mediated oxygen consumption rate changes in mice), but results are variable due to cell type, dose, and context (Malheiro et al., 2023; Puighermanal et al., 2024). Nonetheless, the changes we observed in mitochondria may not solely be due to migration out of the soma. It has been reported that autophagy, including mitophagy, is required for OPC differentiation (Bankston et al., 2019; Yazdankhah et al., 2021). We did not observe any significant effect of BAL cannabis on the presence of autophagosomes in OPCs, but this may be a phenomenon that develops later in the differentiation process (**Supplementary Fig. 5F**).

In summary, OPC differentiation is a highly dynamic and rapid process depending on the context. The changes we observed in OPC morphology and mitochondrial ultrastructure due to BAL cannabis may represent very early stages of OPC differentiation, which we hypothesize could be initiated within the time scale of this study, and warrants further investigation.

### Sex-Dependent Effects

All of the large cannabis-related effects observed in this study were primarily observed in male mice. We cannot claim sex-dependent effects of cannabis type on OPC characteristics in this study as we do not observe significant interactions in our statistical tests (Rich-Edwards and Maney, 2023). However, we do observe large differences in the effect size between male and female mice in specific cannabis groups and are likely underpowered to detect subtle interactions (Morgan, 2025). In line with this, there is a body of literature showing sex effects in oligodendrocyte lineage cells related to cannabis.

RNA-seq data from primary rat cultured OPCs have shown significantly higher expression of *Cnr1* in males compared to females, which could help explain some of the sex-dependent effects in this study (**Supplementary Fig. 1Ai**; Yasuda et al., 2020). However, other datasets, albeit varying by age and region, do not show any strong sex differences in *Cnr1* expression in OPCs (Marques et al., 2016; Heo et al., 2025; Marquardt et al., 2025). That being said, differences in CB1R mRNA have been observed between males and females in rodents in a region-specific manner in the brain (Castelli et al., 2014; Cooper and Craft, 2018; Liu et al., 2020). In addition, numerous studies have shown sex differences in CB1R binding/availability in the rat and human brain, but the direction of the effect is inconsistent (Mateos et al., 2011; Dow-Edwards et al., 2016; Laurikainen et al., 2019; Spindle et al., 2021). At the moment, the literature strongly suggests that there are sex differences in the endocannabinoid system, but findings are too inconsistent and context-dependent to draw any conclusions with respect to its impact on oligodendrocyte lineage cells.

Sex differences in the pharmacokinetics of THC have also been observed (Hložek et al., 2017; Torrens et al., 2020; Baglot et al., 2021; Gazarov et al., 2023; Sallam et al., 2023). However, sex differences are typically linked to differential metabolism of cannabinoids observed after oral consumption or injection, and are rare in studies using inhalation as a route of administration. One study using smoking of cannabis found that the maximum concentration (C_max_) of THC was nearly twice as high in plasma from male C57BL/6J mice—an effect absent in other mouse strains tested—compared to females, but observed little difference in THC levels in the brain (Gazarov et al., 2023). Similarly, little difference was observed in THC or THC metabolites in plasma or brain tissue between sexes following inhalation of THC in other studies (Baglot et al., 2021; Zequeira et al., 2025). In this study, we did not find any strong sex difference in serum THC or 11-OH-THC in any cultivar, but did observe a robust sex difference in serum CBD, supporting the significant interaction between cannabis and sex (Rich-Edwards and Maney, 2023).

A series of publications have investigated the pharmacokinetics of (phyto)cannabinoids after inhalation. Although the authors report that the sample size was too small to make any conclusions about sex differences, vapor from CBD-dominant cannabis and pure synthetic crystalline CBD resulted in comparatively more CBD in whole blood in men compared to women (Spindle et al., 2020; Bergeria et al., 2022). In addition, Sholler et al., (2021) found that women had higher levels of 11-OH-THC in whole blood compared to men immediately following inhalation of vapor from THC-dominant cannabis. These studies hint at sex differences in CBD absorption after inhalation of cannabis vapor and also indicate sex differences in human metabolism of THC, in line with rodent studies. The fact that we observed elevated serum levels of CBD in female mice compared to males exposed to CBD cannabis also suggests sex differences in CBD absorption. However, the direction of the effect is opposite to the studies in humans (Spindle et al., 2020; Bergeria et al., 2022), possibly driven by differing CBD concentrations/cannabis cultivars, exposure paradigm, and species differences. Indeed, when looking at the BAL data, male mice do indeed have higher serum levels of CBD compared to females, in line with the previous human data.

Although not significant, the lower levels of serum 11-OH-THC we observed in female mice compared to male mice exposed to BAL cannabis may also hint at sex differences in THC:CBD metabolism. This could in part be due to relatively less serum THC being present in female BAL mice. Nonetheless, male mice exposed to BAL cannabis had relatively elevated levels of all three of THC, 11-OH-THC, and CBD compared to other cultivars, and in relation to female BAL mice. This ‘triple hit’ may explain why we see marked effects in OPCs only in males exposed to BAL cannabis.

Overall, the large effect caused by BAL cannabis on OPCs in only male mice hints at either differential absorption/metabolism of inhaled phytocannabinoids, sex-specific differences in the OPC response to phytocannabinoids, or a combination of both. Due to a lack of sex difference data in the literature surrounding the endocannabinoid system in oligodendrocyte lineage cells, as well as the response to different cannabis cultivars, explanations remain speculative. This highlights the need for increased research on the sex differences of oligodendrocyte lineage cells and the endocannabinoid system, and their response to (phyto)cannabinoids

### Cultivar-Dependent Effects

Intriguingly, many of the significant cannabis-related findings are related to the BAL cannabis cultivar. We utilized inhalation as the route of administration, which avoids first-pass metabolism of phytocannabinoids in the liver, where many of the influential effects of CBD on THC metabolism are thought to occur (Bornheim et al., 1995; Hložek et al., 2017; Chayasirisobhon, 2020; Zamarripa et al., 2023; Chester et al., 2026). The few studies that have looked at the impact of cannabinoids on OPCs in rodents have primarily used injection of THC (or CBD) alone, and significant effects have been observed (Mecha et al., 2012; Aguado et al., 2021; Huerga-Gómez et al., 2021; Sánchez-de la Torre et al., 2022). However, we observed little effect of high THC or high CBD cannabis cultivars on OPCs. This could be partially due to the reduced bioavailability of THC and CBD from inhalation *vs.* injection or be a product of the ‘entourage effect’ from full-spectrum cannabis flower versus purified alternatives, although data with respect to this effect is minimal (Hložek et al., 2017; Meyer et al., 2018; Chayasirisobhon, 2020; Baglot et al., 2021; Christensen et al., 2023). Another possibility is that the effects of THC alone on OPCs require a longer time scale to be fully appreciated.

Alternatively, we hypothesize that the BAL cannabis cultivar has such a noticeable effect on OPCs because it may act on multiple different pathways that simultaneously drive OPC function in a similar direction. As discussed, THC has been shown to increase OPC differentiation and maturation through the CB1R in mice (Sánchez-de la Torre et al., 2022). In addition, adenosine has been shown to promote OPC cell cycle arrest and subsequent differentiation in rodents (Stevens et al., 2002; Coppi et al., 2015). CBD has been shown to increase extracellular levels of adenosine through inhibition of ENT-1, which is responsible for adenosine uptake (Carrier et al., 2006). CBD has also been shown to act as a FAAH inhibitor, thereby inhibiting degradation of endocannabinoids (i.e., AEA) and endocannabinoid-like lipids (i.e., oleoylethanolamide and palmitoylethanolamide) that act on PPARγ (De Petrocellis et al., 2011; Panlilio et al., 2013). However, this remains to be examined thoroughly in oligodendrocytes. Nonetheless, oligodendrocyte lineage cells have the highest expression levels of enzymes responsible for the formation (*nape-pld*) and degradation (*Faah*) of AEA compared to other cell types in the mouse brain, showing a graded increase as the lineage progresses (**Supplementary Fig. 1**; Zhang et al., 2014). Despite this, the function and importance of these enzymes or of AEA in the oligodendrocyte lineage are relatively unknown. Both THC and AEA are partial agonists of the CB1R with similar efficacy, although THC has stronger binding affinity (Hempel and Xi, 2022). In addition, 11-OH-THC has been shown to be more potent than THC in terms of intoxication, and has a similar binding affinity for CB1R to THC (Lemberger et al., 1972; Wiley et al., 2021; Zagzoog et al., 2022, 2024). If AEA levels do increase due to BAL cannabis and 11-OH-THC has direct effects on oligodendrocyte lineage cells that are comparable to THC, the combined actions of AEA, THC, and 11-OH-THC could exert greater influence on OPC differentiation than THC alone.

Another possible explanation for the effect observed due to BAL cannabis compared to the other cultivars may be related to outcomes on neuronal activity. Not only does THC influence the balance between excitation and inhibition through activation of CB1R, but CBD also alters this balance and network connectivity more broadly through its actions on TRPV1, GPR55, 5-HT1A, and A_2A_ in mice and humans (Pretzsch et al., 2019; Schouten et al., 2024). Callosal OPCs form direct synaptic contacts with both excitatory and inhibitory neurons (Bergles et al., 2000; De Biase et al., 2010; Mount et al., 2019), and neuronal activity influences OPC differentiation, cell cycle progression, proliferation, migration, and expression profiles in mice (Barres and Raff, 1993; de Faria Jr. et al., 2019; Moura et al., 2023). In this way, this two-pronged alteration from THC and CBD on neuronal activity caused by BAL cannabis may indirectly initiate the early phases of OPC differentiation to a greater extent than THC or CBD alone.

Overall, BAL cannabis likely has both direct and indirect effects on OPCs from multiple—perhaps synergistic—mechanisms that result in acute changes to OPCs reminiscent of the early stages of differentiation. Whether or not these OPCs complete differentiation, and what proportion of the newly-formed oligodendrocytes survive to be integrated into the oligodendrocyte pool, is unknown. The integration of newly differentiated OPCs is uncommon, with the majority of newly-formed oligodendrocytes undergoing apoptosis relatively soon after differentiation in adult mice (Hughes et al., 2018; Kamen et al., 2025). If these changes aren’t persistent, how long does it take for these acute changes to revert back to baseline once (phyto)cannabinoids are cleared from the system? These questions should be addressed in future studies.

## Conclusion

In this study, adult male and female mice were exposed to different cultivars of vaporized cannabis to investigate the acute impact of differing ratios of THC:CBD on oligodendrocyte lineage cells in the forceps minor 30 minutes post-onset of cannabis inhalation. We observed changes to OPC morphology and mitochondrial dynamics with a large effect size in response to vaporized cannabis balanced in THC and CBD in adult male, but to a lesser extent in female mice. These observations are reminiscent of the changes observed during OPC differentiation. These data show the differential impact of different cultivars of cannabis and the rapidity with which OPCs respond to phytocannabinoids. Future research should examine the fate of these OPCs and the overall consequences on the oligodendrocyte pool, and in particular what these changes may mean for adaptive myelination, neurocircuitry, and possible effects on behavior. Lastly, this work indicates that more research focusing on sex differences of the endocannabinoid system in oligodendrocyte lineage cells and the effect of phytocannabinoid metabolites is needed.

## Acknowledgments

We acknowledge and respect the ləkʷəŋən (Songhees and X^w^sepsəm/Esquimalt) Peoples on whose territory the University of Victoria stands, and the ləkʷəŋən and WSÁNEĆ Peoples whose historical relationships with the land continue to this day. We acknowledge that image analysis for this work was performed in the University of British Columbia Life Sciences Institute Imaging Core Facility, RRID: SCR_023783. CJM is funded by a Canadian Institutes of Health Research (CIHR) Canada Graduate Scholarship-Doctoral, and was previously supported by the CIHR Canada Graduate Scholarship-Masters and a graduate grant from the Branch Out Neurological Foundation. SL was supported by a CIHR Canada Graduate Scholarship-Masters. HAV was supported by a Fellowship from CIHR and a Brain Canada Rising Star—Canadian Consortium for the Investigation of Cannabinoids (CCIC) Neuroscience Fellowship in Cannabis and Cannabinoid Research—and was a Michael Smith Health Research British Columbia (MSHRBC) Research Trainee. The Tremblay Lab’s Zeiss Crossbeam 350 scanning electron microscope was acquired with funding from the Canada Foundation for Innovation John R. Evans Leaders Fund grant (39965 Laboratory of ultrastructural insights into the neurobiology of aging and cognition) awarded to MET. MET is a Tier 1 Canada Research Chair in *Neurobiology of Healthy Cognitive Aging*. Experiments in the Tremblay Lab were covered through a start-up grant from the School of Medical Sciences, University of Victoria. HHAT was supported by a Post-doctoral Fellowship from CIHR and a Brain Canada Rising Star—Canadian Consortium for the Investigation of Cannabinoids (CCIC) Neuroscience Fellowship in Cannabis and Cannabinoid Research Award. JYK is supported by a CIHR Tier 2 Canada Research Chair in Translational Neuropsychopharmacology.

## Author Contributions (CRediT)

CJM, HT, HK, HAV, JK, and MET—conceptualization, methodology. CJM, HT, HK, EO, SF, SL, MR, and HAV—Investigation. CJM—Formal Analysis, software, visualization, writing (original draft), and writing (review and editing). JK and MET—Resources. HHAT, HAV, JK, and MET—Supervision, funding acquisition, and writing (review and editing). CJM, HAV, JK, and MET—Project management.

## Data Availability

The data that support the findings of this study are available from the corresponding author upon reasonable request.

## Conflicts of interest

The authors declare no conflicts of interest.

## Figures

**Supplementary Fig. 1.**
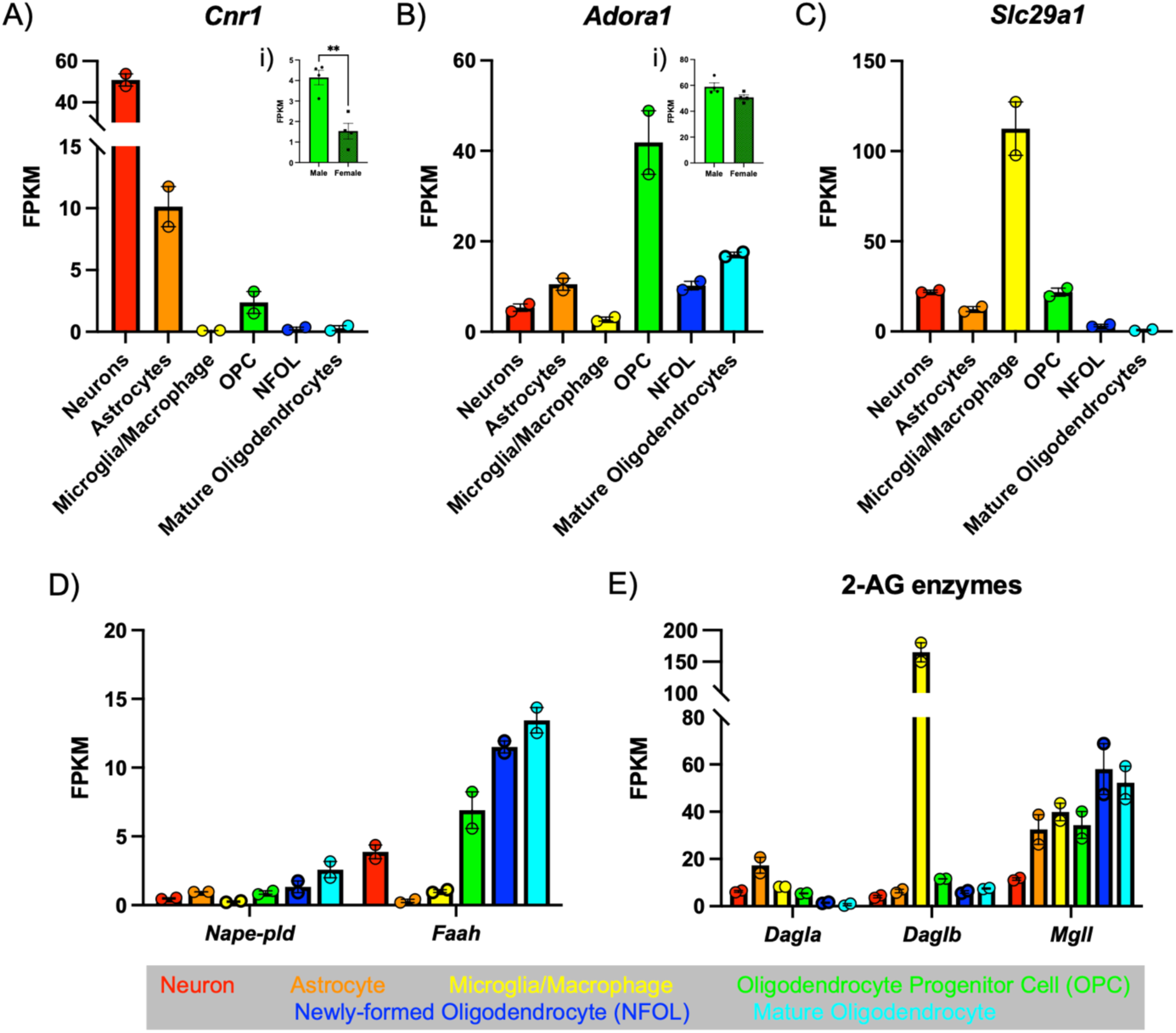
RNAseq expression levels of select genes acquired from publicly available datasets. (A–E) *Cnr1, Adora1, Slc29a1, Nape-pld, Faah, Dagla, Daglb,* and *Mgll* RNA expression levels from the mouse cerebral cortex. Dataset: Zhang *et al.,* (2014). (Ai, Bi) *Cnr1* and *Adora1* RNA expression levels from rat primary cultured OPCs from the cortex of neonatal rats. (Ai) *Cnr1* expression was significantly different (*p-*value = 0.002; *Δ =* 2.617 FPKM; 95% CI 1.341, 3.893) between male and female cultured OPCs, as quantified with an unpaired T-test. Dataset: Yasuda *et al.,* (2020). *Cnr1=*cannabinoid receptor 1, *Adora1*=adenosine A_1_ receptor, *Slc29a1=*equilibrative nucleoside transporter 1, AEA=anandamide, 2-AG=2-arachidonoylglycerol, *Nape-pld*=N-acylphosphatidylethanolamine phospholipase D/synthesizing enzyme for anandamide, *Faah*=fatty acid amide hydrolase/degrading enzyme for AEA, *Dagla/Daglb*=diacylglycerol lipase/synthesizing enzymes for 2-arachidonoylglycerol, and *Mgll*=monoacylglycerol lipase/degrading enzyme for 2-arachidonoylglycerol. FPKM=Fragments Per Kilobase of transcript per Million mapped fragments.

**Supplementary Fig. 2.**
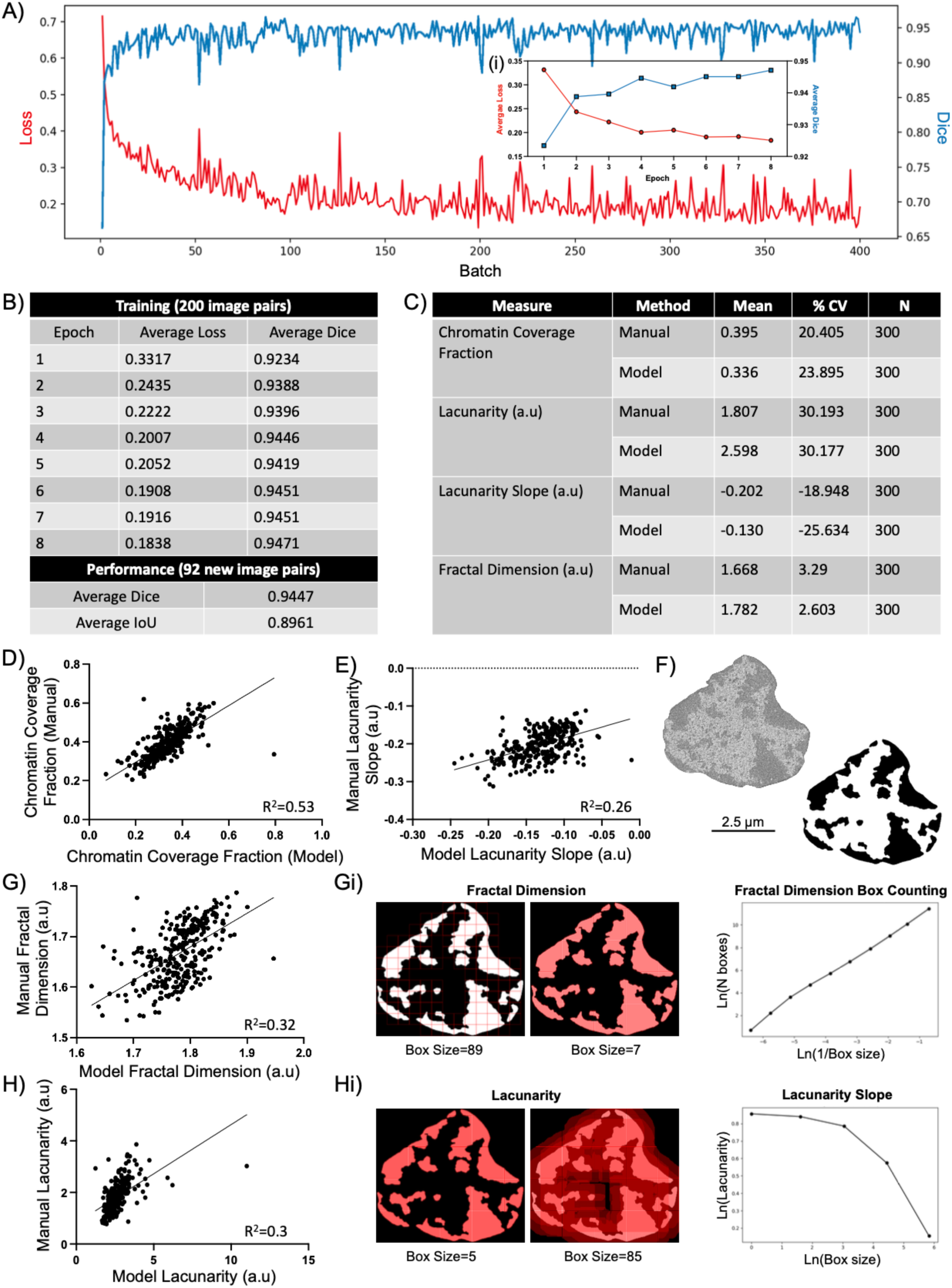
Training performance of automated U-Net segmentation of nuclei and comparison against manual segmentation. (A) Training performance of the U-Net segmentation model with training Loss and Dice plotted against each training batch—which each contains a different subset of training images—and (Ai) the average Loss and Dice per epoch. (B) The average Loss and Dice values per epoch are shown for the same 200 image pairs, as well as the model’s performance in terms of Average Dice and Average IoU on 92 new image pairs that the model was not trained on. (C) Comparison between the manual and U-Net model for chromatin coverage fraction, lacunarity, lacunarity slope, and fractal dimension in terms of mean and % CV. Simple linear regressions are shown for (D) chromatin coverage fraction, (E) lacunarity slope, (G) fractal dimension, and (H) lacunarity for the manual *vs* U-Net model with corresponding R^2^ values. Example images of the U-Net chromatin segmentation are shown in (F), as well as the fractal dimension (Gi) and lacunarity (Hi) with corresponding slopes. % CV=Percent Coefficient of variation, IoU=Intersection over union.

**Supplementary Fig. 3.**
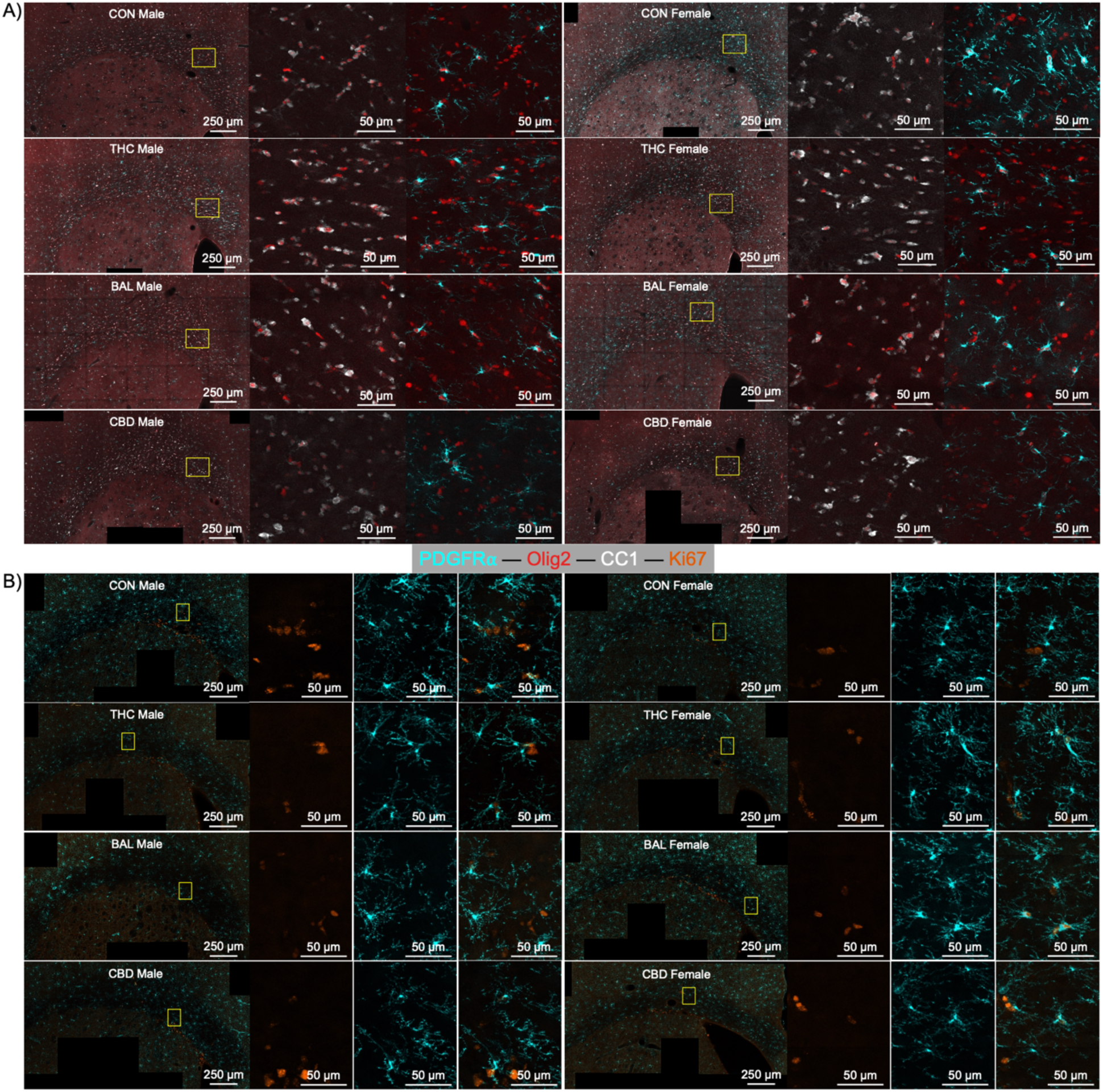
Representative immunofluorescent images for each group for (A) density and distribution data of oligodendrocyte lineage cells, and (B) markers of proliferation in OPCs. Each group has expanded insets to show greater detail of immunofluorescent staining, and the region they were acquired from is depicted by a yellow box on the original image. CON=control; THC=high THC cultivar; BAL=balanced cultivar; CBD=high CBD cultivar.

**Supplementary Fig. 4.**
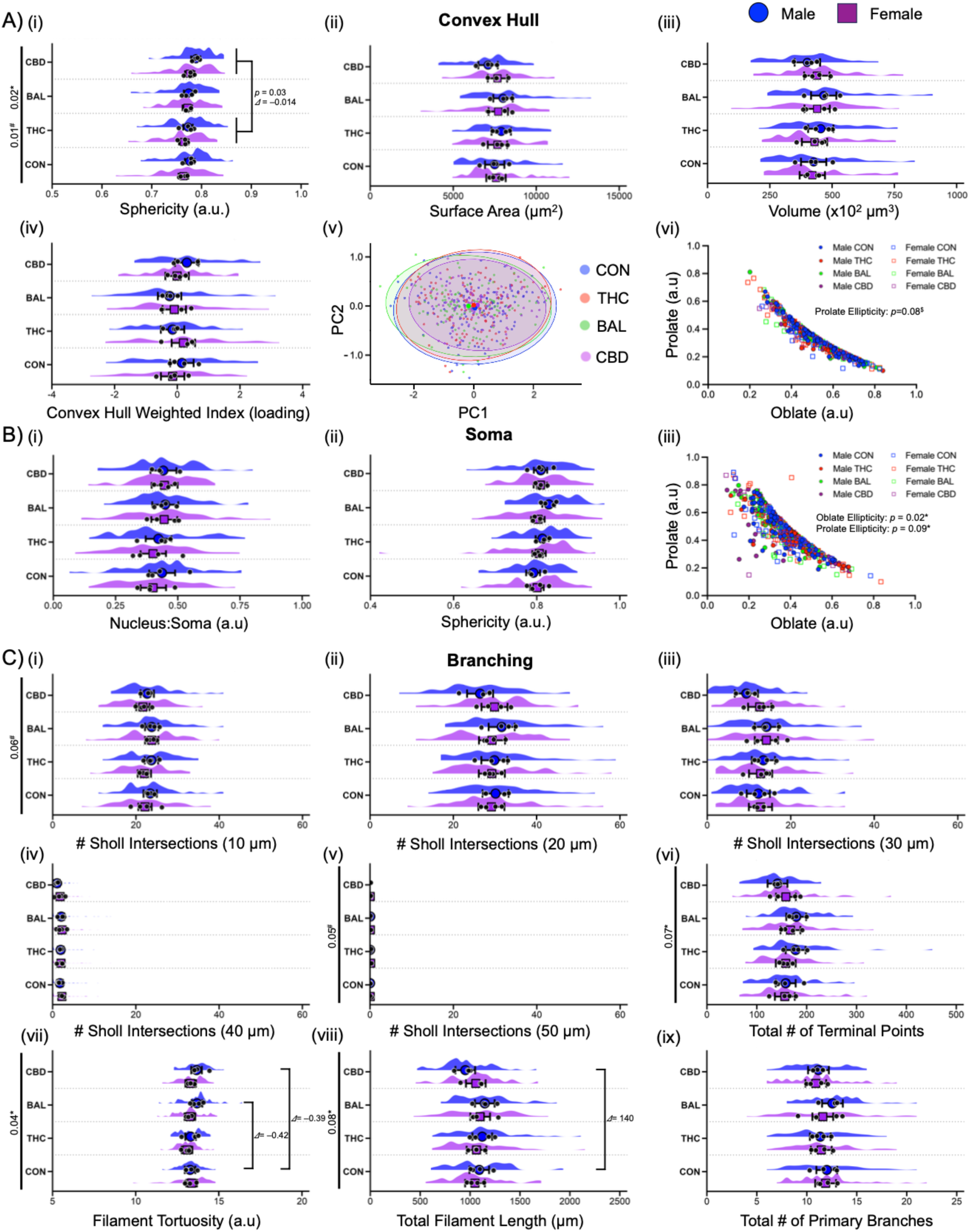
Measures related to the morphology of OPCs. (A) Convex hull measurements: (i) sphericity, (ii) surface area, (iii) volume, (iv) morphological index, (v) principal component 1 vs 2, and (vi) prolate *vs* oblate eccentricity. No significant differences were observed for convex hull, with the exception of a main effect of cannabis (*p*-value = 0.02) and sex (*p*-value = 0.01) for sphericity. (B) Soma measurement: (i) nucleus to soma ratio, (ii) sphericity, and (iii) prolate *vs* oblate eccentricity. No significant differences were observed for these soma measures. (C) Branching measurements: (i) number of Sholl intersections at (i) 10 µm radius, (ii) 20 µm radius, (iii) 30 µm radius, (iv) 40 µm radius, (v) and 50 µm radius, (vi) total number of terminal branch points, (vii) filament/branch tortuosity, (viii) total filament/branch length, (ix) and total number of primary branches. No significant differences were observed for these branching measures, with the exception of a main effect of cannabis (*p-*value = 0.04) for filament tortuosity. Linear-mixed effects model (Aii–iv, Ci–ix) or generalized linear-mixed model with beta family distribution (Ai, vi, Bi–iii) with data expressed as group model-based mean (blue circle or purple square) ± 95% confidence intervals, alongside raw animal means (black circles) and the spread of raw data from each analyzed OPC. Šídák *post-hoc* applied to correct for multiple comparisons, where appropriate. N=4 animals/group/sex, 15 cells/animal. Blue circles=Male; Purple squares=female. * corresponds to effect of cannabis, # corresponds to effect of sex, and $ corresponds to an interaction. *p*-values shown alongside differences in group means (*Δ*) and Cohen’s D (*d*) for comparisons of interest. CON=control; THC=high THC cultivar; BAL=balanced cultivar; CBD=high CBD cultivar.

**Supplementary Fig. 5.**
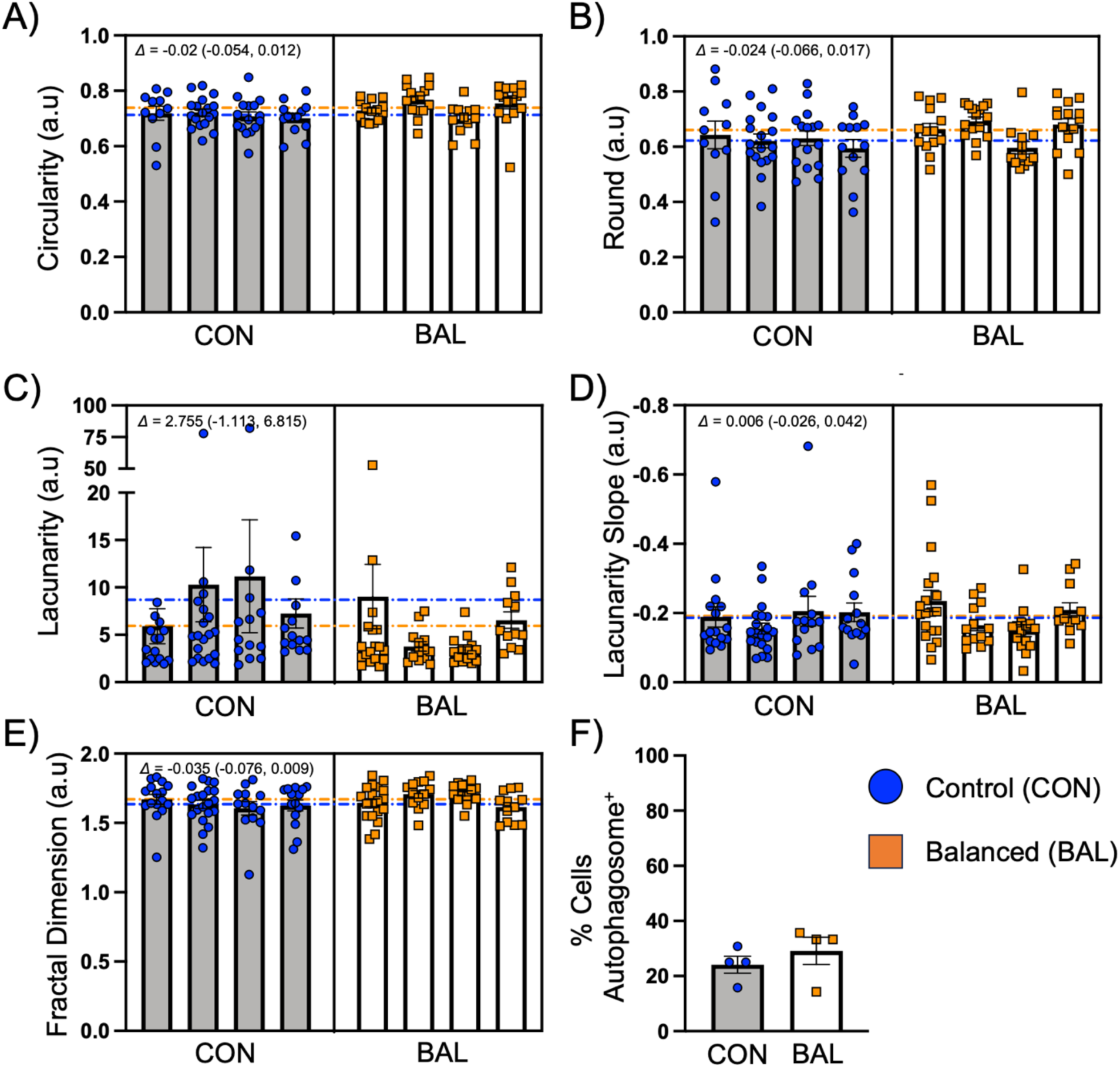
Measures related to the ultrastructural analysis of OPCs. We observed no significant effect of cannabis on (A) mitochondrial circularity or (B) roundness. We also observed no significant effect of cannabis on heterochromatin distribution, including (C) lacunarity, (D) lacunarity slope, or (E) fractal dimension. Lastly, we observed no significant effect of cannabis on the (f) percentage of OPCs with autophagosomes. Linear-mixed effects model (C–E), generalized linear-mixed effects model with beta family distribution (A, B), or unpaired T-test (F). For mitochondria data, mitochondria are nested within cell, and cell nested within group. Each bar represents one animal, and data points represent the average value for each cell analyzed, with the exception of (F), where each point represents an animal and bars correspond to group means. Data expressed as mean ± standard error of the mean. Dotted lines correspond to raw group averages. N=4 animals/group/sex, 11–20 cells/animal. Blue circles=control male; Orange squares=balanced cannabis males. Differences between group means (*Δ*) shown alongside 95% confidence intervals. CON=control; BAL=balanced cultivar.

**Supplementary Fig. 6.**
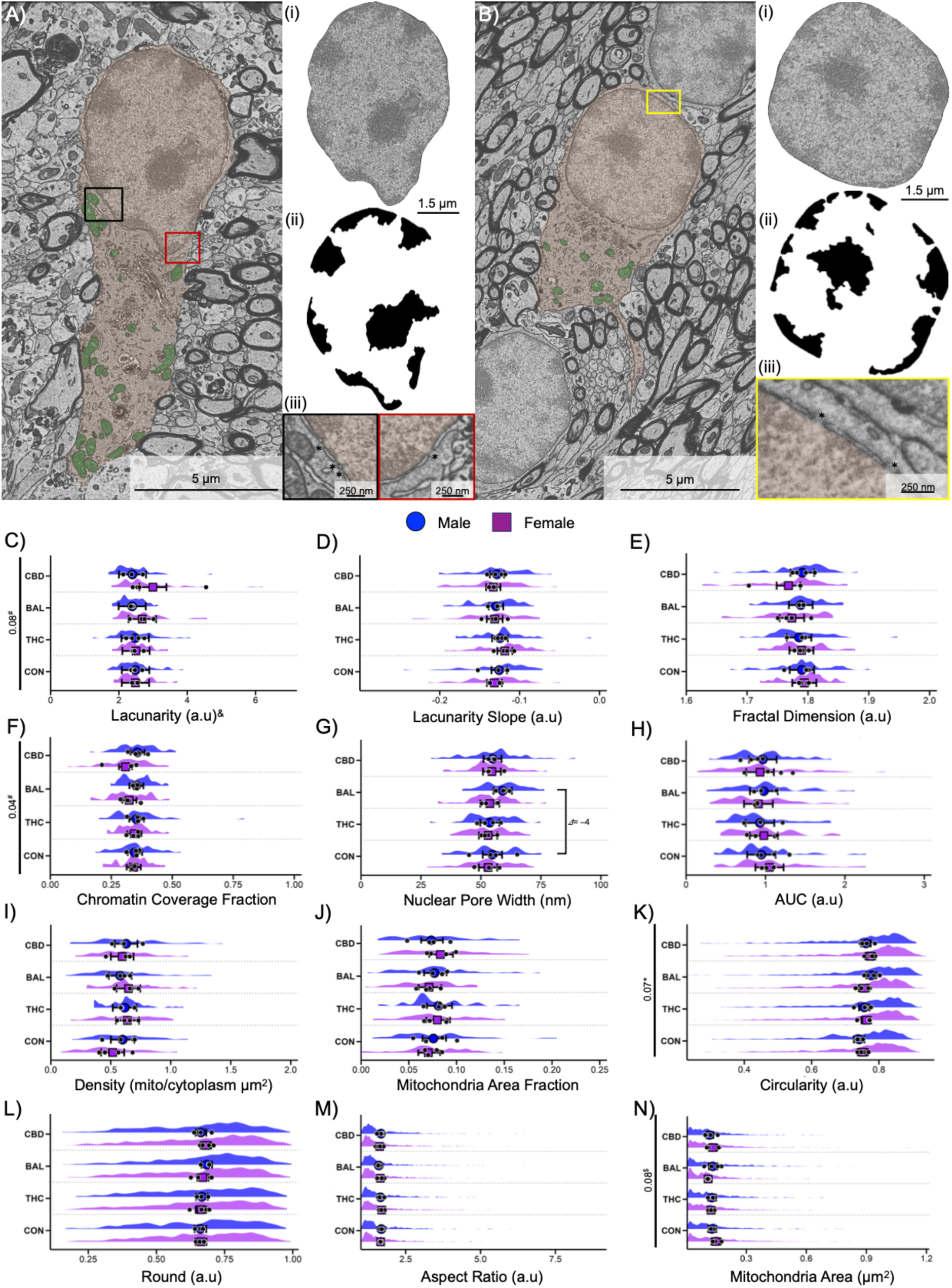
Acute inhalation of vapor from different cannabis cultivars does not alter mature oligodendrocyte ultrastructural characteristics. (A, B) Representative images of two mature oligodendrocytes, including (Ai, Bi) oligodendrocyte nuclei, (Aii, Bii) their heterochromatin segmentation, and (Aiii, Biii) examples of nuclear pores. (A, B) Oligodendrocytes are shaded in orange and mitochondria are shaded in green. (C–E) We observed no significant effect of cannabis on heterochromatin distribution, or (F, H) coverage. We also observed no significant effect on (G) nuclear pore width. Similarly, we saw no significant effect of cannabis on (I) mitochondria density, (J) area fraction, (K) circularity, (L) roundness, (M) aspect ratio, or (N) area. Linear-mixed effects model (C–E, G–I, M, N) or generalized linear-mixed model with beta family distribution (F, J–L) with data expressed as group model-based mean (blue circle or purple square) ± 95% confidence intervals, alongside raw animal means (black circles) and the spread of raw data from each analyzed oligodendrocyte. N=4 animals/group/sex, 10–20 cells/animal. Blue circles=Male; Purple squares=female. * corresponds to effect of cannabis, # corresponds to effect of sex, and $ corresponds to an interaction. *p*-values shown alongside differences in group means (*Δ*) for comparisons of interest. CON=control; THC=high THC cultivar; BAL=balanced cultivar; CBD=high CBD cultivar.

**Supplementary Fig. 7.**
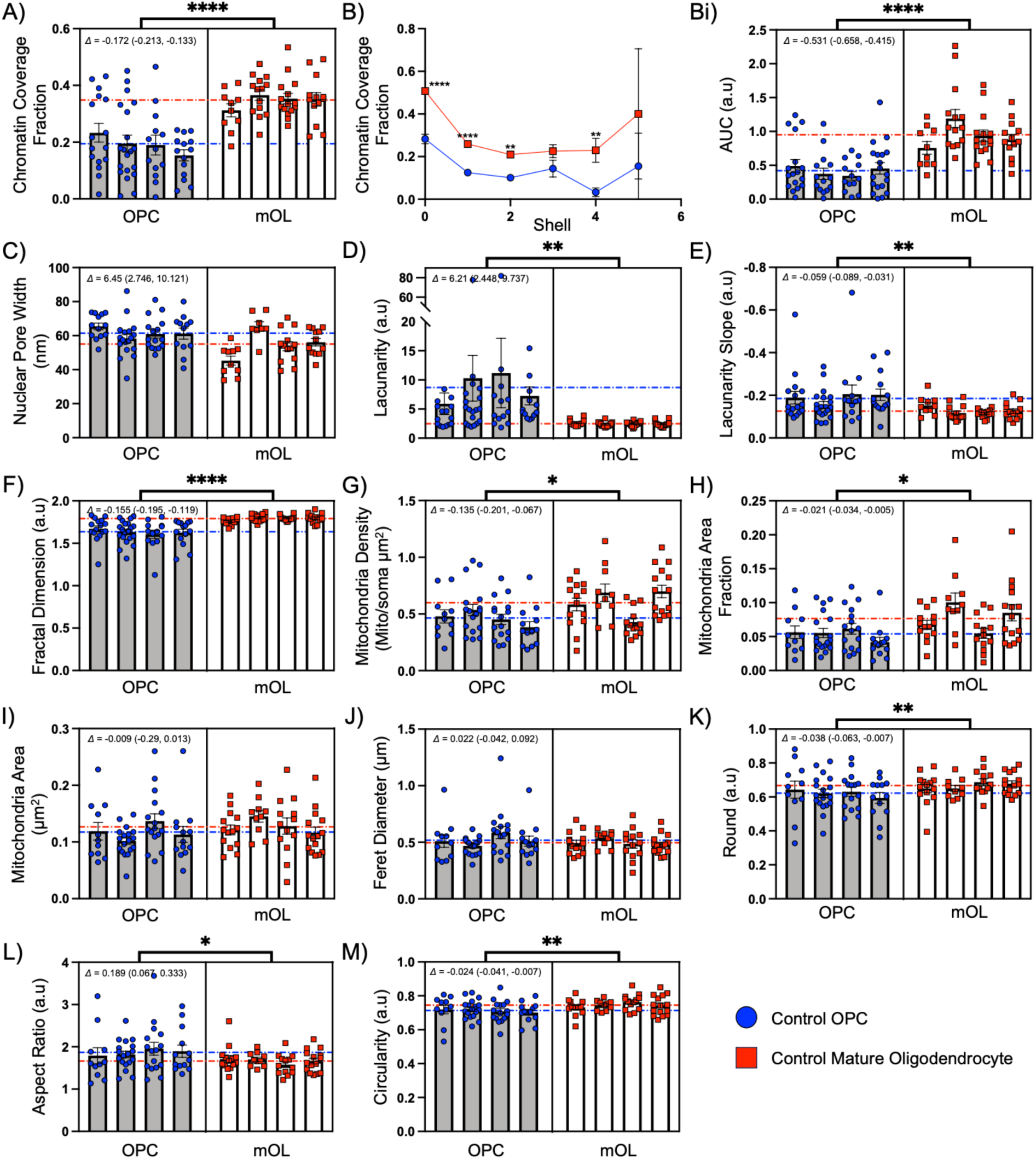
Heterochromatin coverage and distribution, as well as mitochondria shape significantly differ between OPCs and mature oligodendrocytes in control males at an ultrastructural level. (A) The fraction of the nucleus covered by heterochromatin is significantly higher (*p*-value <0.0001) in mature oligodendrocytes compared to OPCs. In addition, the distribution of heterochromatin significantly differs in mature oligodendrocytes compared to OPCs, in terms of (Bi) area under the curve (*p*-value = 0.0001), (D) lacunarity (*p*-value = 0.003), (E) lacunarity slope (*p-*value = 0.002), and (F) fractal dimension (*p*-value <0.0001). No difference was observed for (C) nuclear pore width. The (G) density of mitochondria and (H) fraction of cytoplasm occupied by mitochondria was also significantly higher (*p*-value = 0.05 and *p*-value = 0.02, respectively) in mature oligodendrocytes compared to OPCs. In addition, mitochondria from mature oligodendrocytes had higher (K) roundness (*p*-value = 0.009) and (M) circularity (*p*-value = 0.005), and (L) lower aspect ratio (*p*-value=0.02) compared to mitochondria from OPCs, with no observed differences in (I) area or (J) Feret diameter. Linear-mixed effects model with data expressed as animal mean ± standard error of the mean. For mitochondria data, mitochondria are nested within cell, and cell is nested within group. Each bar represents one animal and data points represent the average value for each cell analyzed. Dotted lines correspond to raw group means. N=4 animals/group, 10–20 cells/animal. Blue circles=control male OPCs; Red squares=control male mature oligodendrocytes. * corresponds to effect of cannabis. *p*-value ≤ 0.05*; *p*-value ≤ 0.01**, *p*-value ≤ 0.0001****. Differences between group means (*Δ*) shown alongside 95% confidence intervals. OPC=oligodendrocyte progenitor cell. mOL=mature oligodendrocyte. AUC=Area under the curve.

## Notes

### Competing Interest Statement

The authors have declared no competing interest.

