## Supplementary File 1 for "Cannabis THC:CBD Composition Affects Oligodendrocyte Progenitor Cell Characteristics Following Acute Cannabis Vapor Inhalation in Adult Male and Female Mice"

**Supplementary Methods File:** Liquid chromatography-tandem mass spectrometry (LC-MS/MS) for serum levels of phytocannabinoids after inhalation of cannabis flower vapor.

The reference standards of THC, CBD, and 11-OH-THC and their deuterated internal standards THC-D3, CBD-D3 and 11-OH-THC-D3 were purchased from Sigma-Aldrich Canada (Oakville, ON, Canada). Captiva enhanced matrix removal lipid (EMR-Lipid) 96-well plate (Agilent, Santa Clara, CA, USA) was used to extract THC and 11-OH THC from the samples. Briefly, 250 μL of acetonitrile (acidified with 1% formic acid) was added to each well, then 50 μL of mouse serum and 20 μL of internal standard solution were added. After the sample passed through under positive pressure at 3 psi, the extraction plate was washed with 150 μL of a mixture of water/acetonitrile (1:4, v:v) solution. The effluent was evaporated under nitrogen at 40 °C, and the residual was reconstituted with the mobile phase for subsequent LC-MS/MS analysis. Calibration standards (2–1000 ng/mL) and quality controls (3 ng/mL and 800 ng/mL) were prepared on the day of analysis by spiking standard working solutions into blank mouse serum. The liquid chromatography separation was achieved on a Vanquish Flex UHPLC system (Thermo Scientific, Waltham, MA, USA). Five microliters of serum extracts were injected and separated on an ACQUITY UPLC BEH C18 Column (1.7 µm, 2.1 mm × 50 mm; Wexford, Waters, Ireland) connected with a VanGuard UPLC BEH C18 Pre-Column (Wexford, Waters, Ireland). The auto sampler was kept at 4 °C and column temperature was at 35 °C. The mobile phase consisted of the following. A: 10 mM ammonium formate with 0.1% formic acid aqueous solution, and B: acetonitrile with 0.1% formic acid. The flow rate was 400 μL/min under a gradient mode. The gradient conditions were sustained as follows: mobile phase B linearly ramped up from 40% to 95% from 0.1 to 4 min, and maintained at 95% for 2 min, then ramped back to 40%. THC, CBD and 11-OH-THC were eluted at 4.6, 4.0, and 3.5 min, respectively, with a total run time of 7 min. MS analysis was conducted with a Q Exactive Focus Orbitrap mass spectrometer (Thermo Scientific, Waltham, MA, USA) equipped with an Ion Max source in positive electrospray ionization (ESI) mode. The source conditions were optimized as the spray voltage of 3.5 kV, the capillary temperature of 300 °C, and aux gas heater temperature of 425 °C. Data were acquired and processed in parallel-reaction monitoring (PRM) mode using TraceFinder™ software (v. 4.1, Thermo Scientific, Waltham, MA, USA). In this PRM mode, protonated 11-OH-Δ9-THC ion (m/z 331.23), CBD ion (m/z 315.23), and Δ9-THC ions (m/z 315.23) were selected as precursors, then fragmented in the higher-energy C-trap dissociation (HCD) cell at collision energy of 20 eV for 11-OH-THC, and 25 eV for THC and CBD. The resulting MS/MS product ions were detected in the Orbitrap at a resolution of 17,500 (FWHM at m/z of 200) with AGC target set at 1 × 10^5^. The most abundant fragments from the MS/MS spectra (m/z 313.22 for 11-OH-THC, and m/z 193.12 for THC and CBD) were selected as the quantifying ions. Other specific fragments, m/z 193.12 for 11-OH-THC, and m/z 259.17 for THC and CBD, were selected as the confirming ions. The resulting chromatograms were extracted and reconstructed with a mass accuracy of 5 ppm for quantification and confirmation. The optimized MS/MS compound parameters are summarized in **Table 1**.

**Table 1.** Optimized LC-MS/MS compound parameters for quantitation of 11-OH-THC, CBD, and THC using PRM mode (CE: collision energy, *m*/*z*: mass/charge ratio, RT: retention time).

| **Analyte and Internal Standard** | **Precursor Ion (*m*/*z)*** | **CE** | **Quantitation Ion (*m*/*z)*** | **Confirming Ion (*m*/*z)*** | **RT (min)** |
| --- | --- | --- | --- | --- | --- |
| 11-OH-Δ9-THC | 331.23 | 20 | 313.22 | 193.12 | 3.5 |
| 11-OH-Δ9-THC–D3 | 334.24 | 20 | 316.23 | 196.14 | 3.5 |
| CBD | 315.23 | 25 | 193.12 | 259.17 | 4.0 |
| CBD-D3 | 318.25 | 25 | 196.14 | 262.19 | 4.0 |
| Δ9-THC | 315.23 | 25 | 193.12 | 259.17 | 4.6 |
| Δ9-THC–D3 | 318.25 | 25 | 196.14 | 262.19 | 4.6 |

CE=collision energy, *m/z*=mass/charge ratio, RT=retention time
